# Downstream-of-gene (DoG) transcripts contribute to an imbalance in the cancer cell transcriptome

**DOI:** 10.1101/2024.01.01.573830

**Authors:** Pedro A. Avila-Lopez, Jessica Xu, Nefertiti Muhammad, Guang-Yu Yang, Shannon M. Lauberth

## Abstract

Downstream-of-gene (DoG) transcripts are an emerging class of noncoding RNAs. However, it remains largely unknown how DoG RNA production is regulated and whether alterations in DoG RNA signatures exist in major cancers. Here, through transcriptomic analyses of matched tumors and non-neoplastic tissues and cancer cell lines, we reveal a comprehensive catalogue of DoG RNA signatures. Through separate lines of evidence, we support the biological importance of DoG RNAs in carcinogenesis. First, we reveal DoG RNAs are tissue-specific and differentially expressed in tumors versus paired normal tissues with their respective host genes involved in tumor promoting versus tumor suppressor pathways. Second, increased DoG RNA number and length is associated with poor patient prognosis. Third, depletion of essential enzyme Topoisomerase I in colon cancer alters RNA polymerase II chromatin engagement leading to termination defects and induction of DoG RNAs. Our results underlie the significance of DoG RNAs in diversifying the cancer transcriptome.

## Introduction

Molecular discoveries revealing alterations in the cancer cell transcriptome have advanced our ability to predict disease severity and understand how cancer cells maintain their proliferative potential, evade tumor suppression and cell death, and undergo cancer cell invasion and metastasis (*1*). Long ncRNAs (lncRNAs) are emerging as key regulators of a variety of cellular processes that influence disease states including cancer (*2*). Yet, the common and specific signatures of lncRNAs across human cancers and the mechanisms driving alterations in their expression in carcinogenesis remain to be explored. An emerging class of lncRNAs referred to as downstream-of-gene (DoG)- containing transcripts (*3, 4*) are initiated at the promoter of upstream protein coding genes in response to stress stimuli that include viral infection (*5, 6*), heat shock (*7*), and osmotic stress (*8*). Precisely, how stress signals trigger DoG RNA production is not fully understood. However, recent evidence has linked DoG biogenesis to defects in transcriptional termination (*8, 9*). Moreover, while these lncRNAs are emerging as products of readthrough transcription in response to stress stimuli, their identity and classification in normal tissues and tumorigenesis remain to be explored.

RNA Polymerase II (RNAPII) is highly processive and termination mechanisms ensure RNAPII comes to a proper halt at protein coding gene ends. Previous studies have established a model for coordination between pausing of the elongating RNAPII and recruitment of the 3’ end processing factors downstream of the poly(A) sites on human genes (*10, 11*). Recent studies provide support for Integrator in targeting promoter-proximally paused RNAPII for termination that prevents elongation (*12*) and the induction of readthrough transcription (*8, 9*). More recently, Integrator has also been shown to support the elongation rate of RNAPII and render paused RNAPII into productive RNA synthesis (*13*). Specifically, hyperosmotic stress disrupts Integrator associations with RNAPII, which decreases Integrator binding to DNA and prompts enrichment of stress-induced DoG RNA production (*8*). Despite these advances, the regulatory processes underlying alterations in RNAPII pause release frequency, termination, and DoG RNA production remain to be elucidated.

Essential enzyme, TOP1 supports RNAPII-dependent transcription through its contributions to preinitiation complex formation (*14–20*) and transcriptional elongation (*21*). Importantly, TOP1 through its recruitment by TFIID has been shown to regulate PIC assembly through the formation of an active TFIID-TFIIA complex (*15*). TOP1 also acts directly to stimulate nucleosome disassembly and gene expression (*17*). In addition, paused promoters are particularly sensitive to the TOP1-selective inhibitor, Camptothecin (CPT) (*22*), suggesting that TOP1 may regulate RNAPII pausing, a highly regulated step of the transcription cycle (*23*). Consistent with a role for TOP1 in pausing is a study revealing that BRD4 supports RNAPII pause-release by enhancing TOP1 catalytic activity (*21*). While TOP1 is involved in the early steps of the RNAPII transcription cycle, other than TOP1 preventing replication stress at R-loop-enriched transcription termination sites (*24*), the roles of TOP1 in regulating transcriptional termination have not been elucidated. Moreover, TOP1 overexpression has been identified to be a frequent event associated with colorectal cancer (CRC) (*25, 26*). Yet, the significance of TOP1 dysregulation and its impact on aberrant transcriptional control in CRC remain to be largely elucidated.

Importantly, our study reveals distinct DoG RNA signatures in several major cancer types that are associated with an increased risk of mortality. We further identified and classified differentially expressed DoG RNA signatures in human colon tumors versus paired normal colon tissues. We unveiled that tumor-specific DoG RNAs are produced from host genes that are functionally classified in contributing to tumor promoting pathways. Comparatively, DoG RNAs produced in normal tissues and normal colon epithelial cells were found to be produced from host genes involved in normal developmental and tumor suppressor pathways. We also reveal TOP1 as a central regulator of DoG RNA production in colon cancer. Notably, our data uncovered that TOP1 depletion disrupts global RNAPII chromatin engagement by releasing paused RNAPII into productive elongation past the ends of a thousand genes that, in turn robustly support DoG RNA production. Our study provides a comprehensive understanding of dysregulated DoG RNA signatures that have important implications for understanding several major cancer types and promise to elicit new therapeutic targets for tuning gene expression programs that shift disease-related networks.

## Results

### Alterations in DoG RNA expression and length lead to transcriptomic imbalances with clinical relevance in major cancer types

To identify and characterize DoG RNA transcriptional landscapes in four major cancer types, RNA-sequencing (RNA-seq) data obtained from The Cancer Genome Atlas (TCGA) (*27*) was analyzed. Specifically, we analyzed RNA-seq data from a total of 50 patient tissues [25 paired tumor and normal tissues (NTs)] from breast invasive carcinoma (BRCA), colon adenocarcinoma (COAD), liver hepatocellular carcinoma (LIHC), and prostate adenocarcinoma (PRAD). DoG RNA calling was performed using DoGFinder 1. (*28*) and the following criteria that included RNA-seq signal that shows >60% RNA-seq coverage and that is >5 kb of continuous read density downstream of the 3’ end of every annotated gene locus. Compared to paired NTs, breast (n = 2,641), colon (n = 1,365), liver (n = 721), and prostate (n = 2,014), a moderate increase in the number of DoG RNAs was identified in the paired tumors, BRCA (n = 3,164), COAD (n = 3,080), LIHC (n = 1,854), and PRAD (n = 2,269) (Fig. 1A, Table S1). Notably, however, the molecular feature that consistently distinguished DoG RNAs in tumors versus paired NTs across all four tissues was a significantly longer median extension strength, or the continuous RNA-seq signal extending beyond the annotated gene end (Fig. 1A, Table S1). Our definition of DoG RNA length or extension strength is dependent on the defined criteria for DoG calling (>60% RNA seq coverage and >5 kb continuous read density). Thus, these measurements provide relative estimates rather than defined endpoints of DoG RNAs. Compared to paired NTs, breast (11,927 kb), colon (9,577 kb), liver (8,946 kb), and prostate (10,570 kb), the median extension strength in the paired tumors was significantly longer BRCA (13,107 kb), COAD (20,901 kb), LIHC (10,407 kb), and PRAD (12,004 kb) (Fig. 1A, Table S1).

**Fig. 1.**
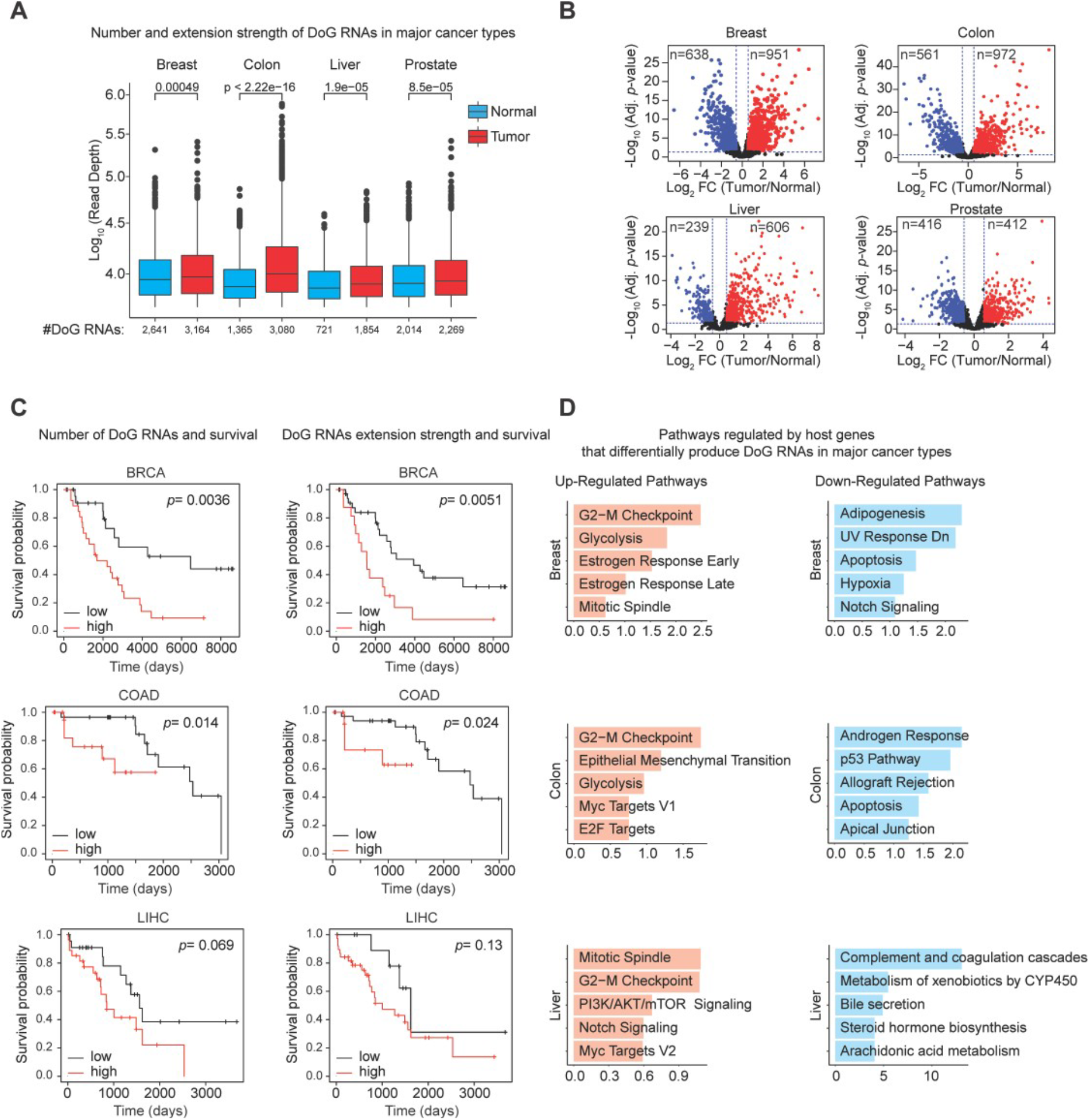
Readthrough transcription is prevalent in cancer. A, Number and extension strength of the DoG RNAs identified by DoGFinder (*28*) in breast, colon, liver, and prostate paired non-neoplastic (n = 25) and tumor (n = 25) samples from TCGA. The extension strength is shown in log10 scale. Boxplots enclose values between first and third quartiles, midlines show medians, and whiskers extend to data points within 1.5 the interquartile range from the box, outliers are shown. Statistical significance was determined by Wilcoxon rank-sum test. *p*-values include 0.00049, <2.22e-16, 1.9e-05, and 8.5e−05 for tumors vs normal, respectively. **B,** Volcano plots of the differentially expressed DoG RNAs (log2 FC>0.58 or log2 FC<-0.58, *q*-value< 0.05) in breast, colon, liver, and prostate cancer compared with normal tissues. The significant down-regulated and up-regulated DoG RNAs are denoted. **C,** Kaplan–Meier curves showing average survival of 50 patients with breast (BRCA), colon (COAD), liver (LIHC) tumors that were examined according to low and high DoG RNA number (right) and DoG RNA extension strength (left). The cutoff were selected by Kaplan-Meier Plotter (*61*). **D,** Top five (*p*-value < 0.05) MSigDB pathways for down-regulated (blue) and up-regulated (red) DoG RNAs in breast, colon, liver, and prostate cancer compared with normal tissues. See also Fig. S1 and Table S1 and S2.

Tumor-specific changes in DoG RNA abundance were determined by measuring the log2-transformed ratio of the signal attributed to DoG transcripts produced from one host DoG-producing gene in a tumor relative to the signal attributed to DoG transcripts of that same gene in a paired NT. DoG RNA expression levels are significantly altered (> or <1.5-fold, *q*-value < 0.05) in the transcriptomes of the tumors versus NTs for all cancer types. Compared to the NTs, we identified a significant number of DoG RNAs that are either upregulated (n = 951, 972, 606, and 412, in BRCA, COAD, LIHC, and PRAD, respectively) or downregulated (n = 638, 561, 239, and 416, in BRCA, COAD, LIHC, and PRAD, respectively) in the respective tumors (Fig. 1B). Comparative analysis of the differentially activated DoG RNAs in BRCA, COAD, LIHC, and PRAD tumors revealed a small number (n = 26) of overlapping DoG RNAs (Fig. S1A). Similarly, only few (n = 3) repressed DoG RNAs were found to overlap among the four tumor types (Fig. S1B). Thus, the vast majority of DoG RNAs that are upregulated (n = 554, 686, 342, and 187, in BRCA, COAD, LIHC, and PRAD, respectively) and downregulated (n = 418, 363, 158, and 212, in BRCA, COAD, LIHC, and PRAD, respectively) are tissue-specific (Fig. S1A,B).

We next integrated the DoG RNA signatures in the BRCA, COAD, LIHC, and PRAD tumors (Fig. 1A) with clinical survival information from 50 patients (Table S2). The prostate cancer patient information was not examined since all of these patients survived (Table S2). The breast, colon, and liver patient information was stratified into two subsets of high versus low median DoG RNA number or extension strength (Fig. 1C, Table S2). As illustrated by the Kaplan-Meier survival plots (Fig. 1C), high versus low DoG number in BRCA and COAD tumors is associated with poor patient survival times with a median survival of 1,692 versus 6,456 days and 899 versus 1,711 days, respectively. In addition, high versus low DoG extension strength is associated with poor patient prognosis and lower survival probabilities in BRCA and COAD cancer patients with median survival of 1,563 versus 3,941 days and 214 versus 1,661 days, respectively (Fig. 1C and Table S2). Comparatively, the survival probabilities were reduced but not significantly for the liver cancer patients with high versus low DoG number and extension strength (Fig. 1C). To further examine the relationship between DoG RNAs and reduced patient survival in colon, breast, and liver cancer patients, we examined functional pathways associated with DoG-producing host genes identified in the BRCA, COAD, and LIHC tumors. Specifically, we found that upregulated DoG RNAs in BRCA, COAD, and LIHC tumors revealed key regulators of tumor promoting pathways, including G2-M checkpoint that was shared among all three tumor types and MYC targets that was shared among COAD and LIHC tumors (Fig. 1D, left). Other tumor promoting pathways identified for either BRCA, COAD, or LIHC tumors included E2F targets and epithelial mesenchymal transition (Fig. 1D, left). In comparison, functional pathway analysis of the DoG-producing host genes associated with downregulated DoG RNAs revealed tumor suppressor pathways (apoptosis, p53 pathway) or normal cellular pathways including metabolism of xenobiotics, steroid hormone biosynthesis, apical junction, and adipogenesis (Fig. 1D, right). Finding that DoG RNAs are produced from genes involved in tumorigenesis is consistent with the finding that increased DoG number and extension strength are associated with poor patient survival. Taken together, these findings are consistent with DoG RNAs serving as a key source of transcriptome diversity that have clinical and biological significance in breast and colon cancer.

## DoG RNAs reshape the transcriptome of human colon tumors

To avoid potential ambiguities associated with differences in experimental execution and analysis of the published datasets, we performed nascent RNA sequencing using rRNA-depleted RNA (total RNA-seq) to identify DoG RNAs in five paired COAD tumors and NTs. Hundreds to thousands of DoG RNAs were identified in the NTs and COAD tumors from each of the patients (Fig. 2A). Relative to the matched NT, a higher DoG number was identified in three of the five COAD tumors, while a comparable number was identified in the other two COAD tumors (Fig. 2A and Table S3). Interestingly, and like the TCGA data analysis (Fig. 1A), DoG extensions were found to be significantly longer in all five COAD tumors with median extension lengths of 22.6 kb and 28.4 kb in the NTs and COAD tumors, respectively (Fig. 2A and Table S3). We also revealed an overlap of 2,006 DoG-producing host genes that were identified in the 5 COAD tumors and the 25 COAD tumors from TCGA (Fig. S2A, left). Specifically, 41% of the DoG-producing host genes in the TCGA COAD tumors show overlap with the host genes identified in the 5 COAD tumors, and conversely, 12% of the host genes in the 5 COAD tumors overlap with those identified in the TCGA datasets (Fig. S2A, left). Similarly, we identified an overlap of 808 DoG-producing host genes in the 5 paired colon NTs and TCGA colon NTs (Fig. S2A, right). Specifically, 35% of the host genes in the TCGA data sets showed overlap with the host genes in the 5 NTs and conversely, 32% of the host genes in the 5 NTs, showed overlap with those in the TCGA datasets (Fig. S2A, right). The comparison of the host genes in the 5 COAD and NTs to the TCGA datasets reveals a lower percentage of overlap that relates to the higher total number (n = 6,350 vs. 3,080 and 6,635 vs. 1,365, respectively) of host genes that we identified using total RNA-seq versus mRNA-seq in the 5 COAD and NTs versus the TCGA datasets (Fig. S2A). Taken together, our data suggests that there exists congruence among the DoG RNAs identified in the TCGA data sets versus our analyses of COAD tumors, showing a consistency in the prevalence of specific DoG RNA signature in cancer.

**Fig. 2.**
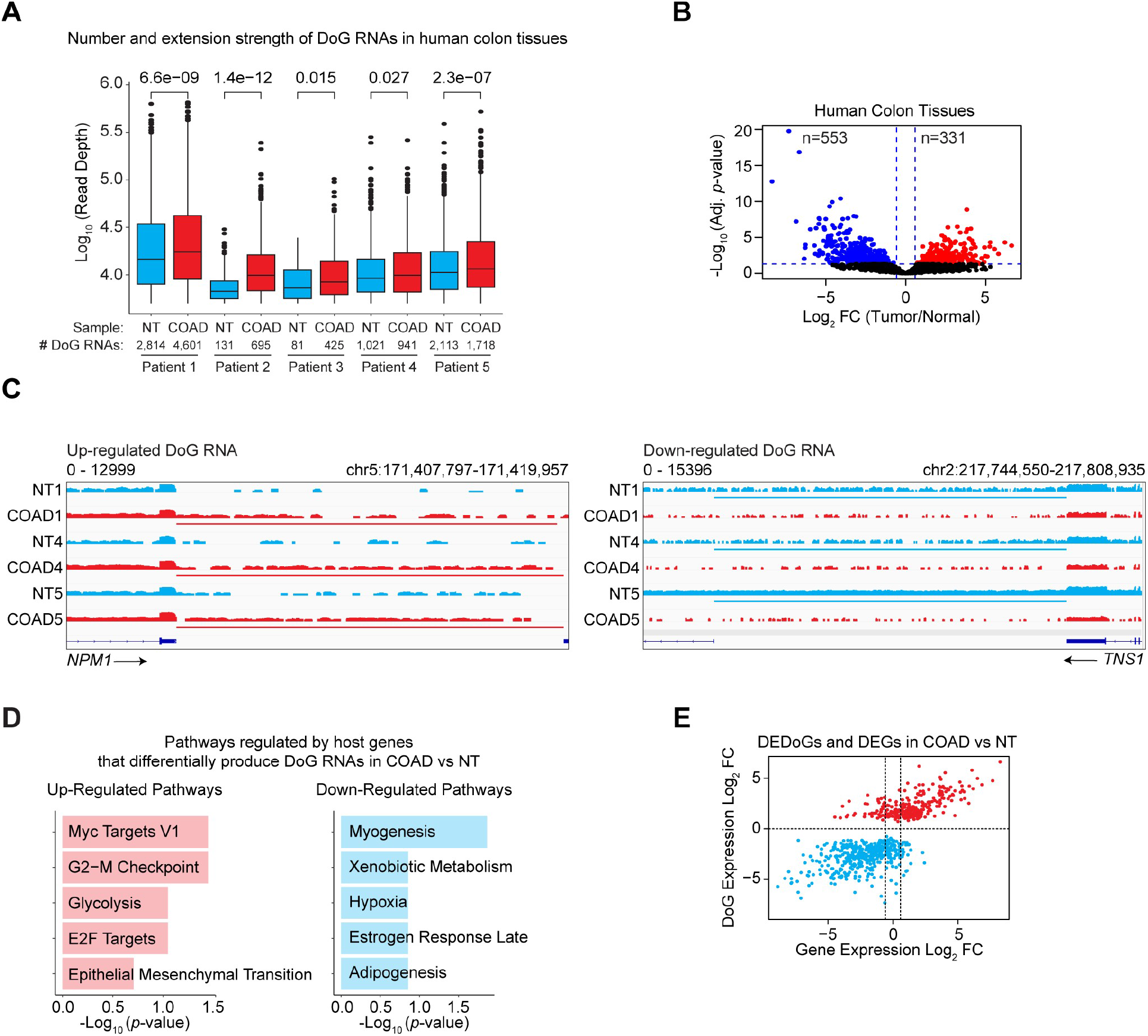
DoG RNAs in colorectal carcinoma tissues are associated with tumorigenic pathways. A, Number and extension strength of DoG RNAs identified by DoGFinder (*28*) in five non- neoplastic (NT) and five colon adenocarcinoma (COAD) tissues. The extension strength is shown in log10 scale. Boxplots enclose values between first and third quartiles, midlines show medians, and whiskers extend to data points within 1.5 the interquartile range from the box, and outliers are shown. Statistical significance was determined by Wilcoxon rank- sum test. *p*-values include 6.6e-09, 1.4e-12, 0.015, 0.027, and 2.3e-07 for patients 1 to 5, respectively. **B** Volcano plot of the differentially expressed DoG RNAs (log2 FC>0.58 or log2 FC<-0.58, *q*-value< 0.05) in five NT and five COAD tissues. The down-regulated and up-regulated significant DoG RNAs are denoted. **C,** IGV tracks of total RNA-seq signal in log RPKM at *NPM1* and *TNS1* loci in NT and COAD samples. NT and COAD tissues are represented in blue and red, respectively. The horizontal bars define the DoG region determined by DoGFinder (*28*). **D,** Top five (*p*-value < 0.05) MSigDB pathways for down-regulated (blue) and up-regulated (red) DoG RNAs in five NT and five COAD tissues. **E,** Scatter plot showing gene Log2 fold change for the DoG-producing gene on the x-axis and the Log2 FC for the DoG RNAs on the y-axis. Down-regulated and up- regulated DoG RNAs in COAD samples relative to NT are represented in blue and red respectively. See also Fig. S2 and Table S3.

Comparative expression analyses of the DoG RNA signatures in the paired NTs versus COAD tumors revealed the expression changes of DoG RNAs that are altered (> or <1.5- fold, *q*-value <0.05) in tumors. Specifically, we found that DoG RNAs were downregulated (n = 553, 6%) or upregulated (n = 331, 4%) in the tumors (Fig. 2B). The percentage of differentially expressed DoG RNAs in tumors is comparable to the percentage of DoG RNAs (∼10%) that were found to be altered in response to hyperosmotic stress (*8*). Differential DoG RNA expression in NTs versus COAD tumors is further evidenced by the increased or decreased RNA-seq signal mapping downstream of the TES of protein coding genes (Fig. S2B). For example, while the *NPM1* gene produced a DoG RNA in COAD tumors versus NTs, the *TNS1* gene produced a DoG RNA specifically in NTs, but not COAD tumors (Fig. 2C). Using RNA purified from four additional COAD tumors and paired NTs, quantitative polymerase chain reaction with reverse transcription (qRT-PCR) revealed that specific DoG RNAs, do*NOLC1* and do*PLD6* are significantly more highly expressed (2.8- to 25-fold and 4- to 80-fold) in the COAD tumors relative to paired NTs, respectively (Fig. S2C). Using the Molecular Signatures Database in Enrichr (*29*), functional pathway analysis of the host DoG-producing genes associated with upregulated DoG RNAs revealed tumor promoting functional pathways including MYC targets, G2-M checkpoint, E2F targets, and epithelial mesenchymal transition (Fig. 2D). Among the pathways associated with the DoG-producing genes with downregulated DoG RNAs are normal cellular processes including myogenesis and xenobiotic metabolism (Fig. 2D). Notably, and consistent with the partial overlap of DoG RNAs that are present in the TCGA COAD and our COAD tumors is the finding that all 5 functional pathways that are tumor promoting (MYC targets, G2-M checkpoint, E2F targets, glycolysis, and epithelial mesenchymal transition) were identified in common for the host genes associated with the upregulated DoG RNAs for both the TCGA (Fig. 1D) and our COAD tumors (Fig. 2D).

We next wanted to determine whether the expression of DoG RNAs relates to the expression levels of their host genes. Log2 fold changes in the expression of DoG RNAs versus DoG-producing host genes in the five COAD tumors versus paired NTs revealed that a large number (n = 219, 66%) of DoG RNAs are produced from transcriptionally active DoG-producing genes (Fig. 2E). In comparison, a smaller number of DoG RNAs are produced from DoG-producing genes whose expression levels are not changing (n = 75, 23%) or have become silenced (n = 37, 11%) in COAD tumors relative to NTs (Fig. 2E). Moreover, downregulated DoG RNAs in COAD tumors relative to NTs are largely produced from DoG-producing genes (n = 416, 75%) that are also downregulated in COAD tumors (Fig. 2E). A smaller number of downregulated DoG RNAs were associated with host genes whose expression levels are not changing (n = 69, 13%) or are upregulated (n = 68,12%) in the COAD tumors (Fig. 2E). These data draw a strong parallel between the transcription levels of DoG RNAs and their respective host genes. Collectively, our observations taken from paired colon tumors and NTs and the TCGA datasets for paired NTs and COAD, BRCA, LIHC, and PRAD tumors support differential expression of DoG RNAs in tumors versus paired NTs that are associated with poor survival probabilities and host genes that are involved in tumor promoting versus tumor suppressor pathways, respectively. This data also suggests a potential function of DoG RNAs in normal context.

## DoG RNAs lead to a length-associated transcriptomic imbalance in colon cancer cell lines

To refine the scope of our observations that there exists differentially expressed DoG RNAs with an extended length in tumors versus paired NTs, total RNA-seq was employed to identify DoG RNAs in colon cancer cell lines, HCT116 and SW480 versus colon epithelial FHC cells. A significant number of DoG RNAs (n = 930, 1,620, and 1,864) exhibiting an average extension strength (13.2, 14.0, and 13.7 kb) were identified in FHC, HCT116, and SW480 cells, respectively (Fig. 3A and Table S4). By comparing differential expression analysis of DoG RNAs in SW480 and HCT116 versus colon FHC cells, we identified significantly upregulated (n = 1,023 or n = 818) and downregulated (n = 1,022 or n = 883) DoG RNAs in SW480 and HCT116 cells, respectively (Fig. 3B). Specifically, heatmaps (Fig. S3A) and specific gene examples, *BUB3* and *EXT1* (Fig. 3C) reveal the upregulated versus downregulated DoG RNAs past the annotated gene ends in HCT116 and SW480 versus FHC cells. To confirm DoG RNA identification in colon cancer SW480 cells, we performed genome-wide mapping of active RNAPII by employing precision nuclear run-on sequencing (PRO-seq) (*30*), which revealed comparable identification of DoG RNAs as observed in our RNA-seq data (Fig. S3B). Moreover, qRT-PCR analyses confirmed differential DoG RNA production by revealing that DoG RNAs (do*NOLC1* and do*PLD6*) produced from host genes, *NOLC1* and *PLD6* are expressed 4- to 15-fold higher in colon cancer cell lines HCT116, SW480, and DLD-1 relative to colon epithelial FHC and HCoEpiC cells (Fig. S3C). Notably, functional pathway analysis of the host genes associated with both up- and downregulated DoG RNAs in HCT116 and SW480 versus FHC cells are strongly correlated with each other and the functional pathways identified for the DoG RNA-producing genes in COAD tumors (Fig. 2D). Specifically, we identified that the host genes associated with upregulated DoG RNAs are linked to pro-tumorigenic pathways including G2-M checkpoint, E2F targets, and MYC targets (Fig. 3D). In comparison, the host genes undergoing a loss of DoG RNA production in HCT116 and SW480 versus FHC cells revealed functional pathways that include the UV response down, epithelial mesenchymal transition, hypoxia, and apical junction pathways (Fig. 3D). Moreover, and consistent with our analysis of DoG RNA signatures in COAD tumors, we found that the expression levels of DoG RNAs are largely consistent with the expression levels of their corresponding DoG RNA-producing host gene (Fig. 3E). Specifically, log2 fold changes in the expression of DoG RNAs versus DoG-producing host genes in both SW480 and HCT116 cells versus FHC cells revealed that the vast majority (n = 862 and 692, 84% and 85%, respectively) of DoG RNAs are produced from transcriptionally active DoG-producing genes (Fig. 3E). In comparison, a smaller number of DoG RNAs are produced from DoG-producing genes that are not changing (n = 104 and 82, 10% and 10%, respectively) or that become silenced (n = 57 and 44, 6% and 5%, respectively) in the cancer cell lines relative to the FHC cells (Fig. 3E). Moreover, a vast majority (n = 591 and 597, 58 and 67%, respectively) of DoG RNAs that are downregulated in SW480 and HCT116 cells relative to FHC cells are largely produced from DoG- producing genes that are also largely downregulated in the cancer cell lines (Fig. 3E). A smaller number of downregulated DoG RNAs were associated with host genes that are not changing (n = 153 and 97, 15% and 11%, respectively) or that are upregulated (n = 278 and 189, 27% and 21%, respectively) in SW480 and HCT116 cells (Fig. 3E). These findings support differential expression of DoG RNAs that are produced in colon cancer cell lines from host genes involved in tumor promoting pathways, which parallels our findings in COAD tumors.

**Fig. 3.**
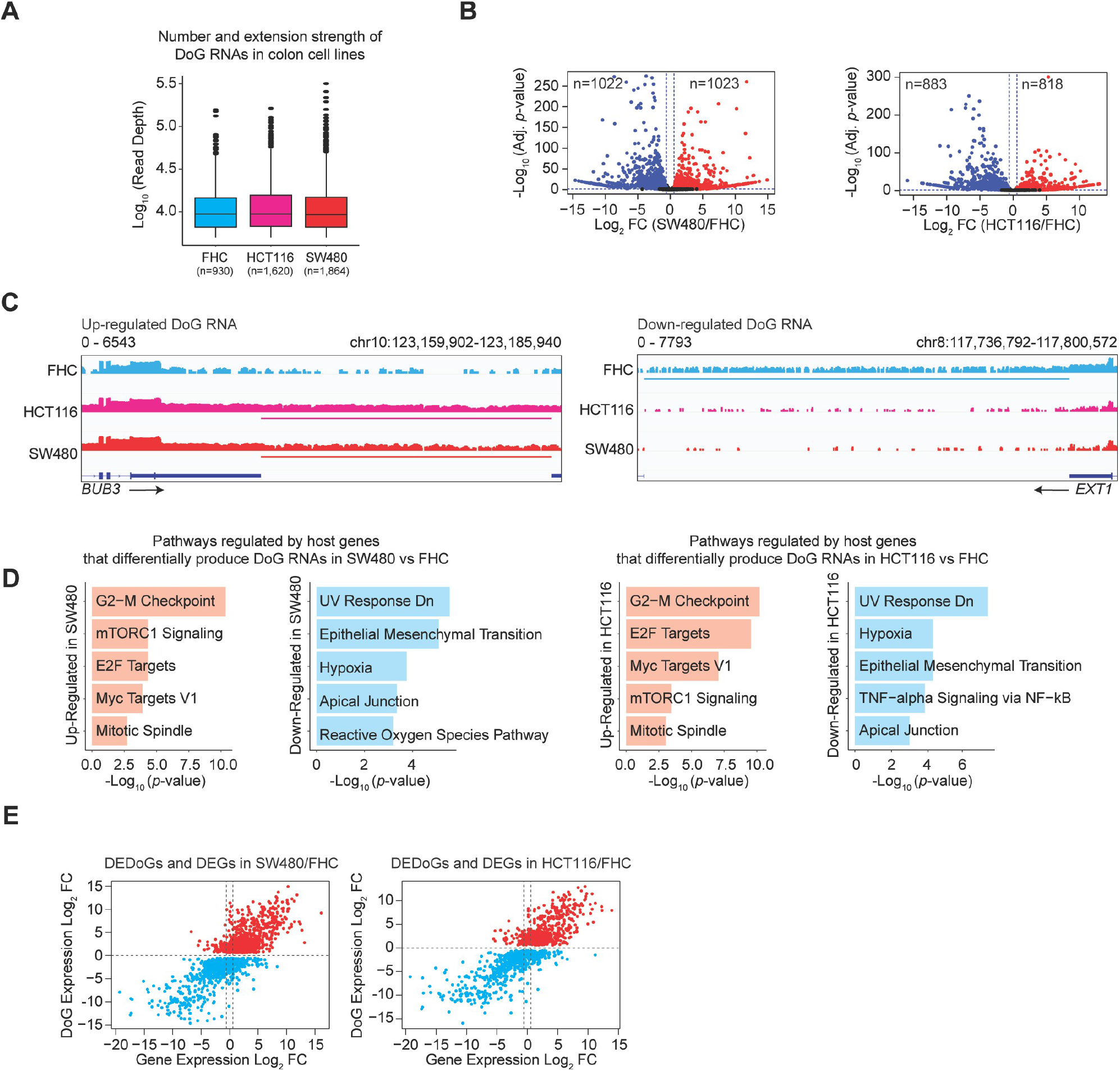
DoG RNA production is prevalent in colorectal carcinoma cell lines. A, Number and extension strength of DoG RNAs identified in FHC, HCT116, and SW480 cells by DoGFinder (*28*). Boxplots enclose values between first and third quartiles, midlines show medians, and whiskers extend to data points within 1.5 the interquartile range from the box, outliers are shown. Statistical significance was determined by Wilcoxon rank-sum test. *p*-value < 0.05. The extension strength is shown in log10 scale. **B,** Volcano plot of the differentially expressed DoG RNAs (log2 FC>0.58 or log2 FC<-0.58, *q*-value< 0.05) in SW480 vs FHC and HCT116 vs FHC cells. The down-regulated and up- regulated significant DoG RNAs are denoted. **C,** IGV tracks of total RNA-seq signal in log RPKM of *BUB3* and *EXT1* loci in FHC, HCT116, and SW480 cells. The horizontal bars define the DoG region determined by DoGFinder (*28*). **D,** Top five (*p*-value < 0.05) MSigDB pathways for down-regulated (blue) and up-regulated (red) DoG RNAs in SW480/FHC (left) and HCT116/FHC (right). **E,** Scatter plots showing gene Log2 fold change for the DoG-producing gene on the x-axis and the Log2 FC for the DoG RNAs on the y-axis. Down-regulated and up-regulated DoG RNAs in SW480/FHC (left) and HCT116/FHC (right) are represented in blue and red respectively. See also Fig. S3 and Table S4.

## TOP1 depletion induced DoG RNA production in colon cancer

To identify potential regulators of DoG RNAs, we sought to identify genes involved in RNAPII-dependent transcription that are differentially expressed in colon cancer. Specifically, we performed differential expression analysis of 161 genes classified according to the GO term, transcription by RNA polymerase II (GO: 0006366) (*31–33*) and by using RNA-seq data from 275 COAD tumors and 349 NTs from TCGA and GTEx databases (*27, 34*). Among the top ten most highly expressed genes in this RNAPII- related subset was *TOP1*, which was identified in COAD tumors as the most highly expressed gene (Fig. 4A). Significantly higher *TOP1* mRNA levels were identified in the 275 COAD tumors versus 349 NTs from TCGA and GTEx databases (*27, 34*) (Fig. S4A). Similarly, as shown in Fig. 4B, immunoblot analysis revealed significantly higher TOP1 protein levels in the five COAD tumors relative to the matched NTs that were used for DoG RNA identification in Fig. 2A. These findings are consistent with the TOP1 dysregulation in colon cancer.

**Fig. 4.**
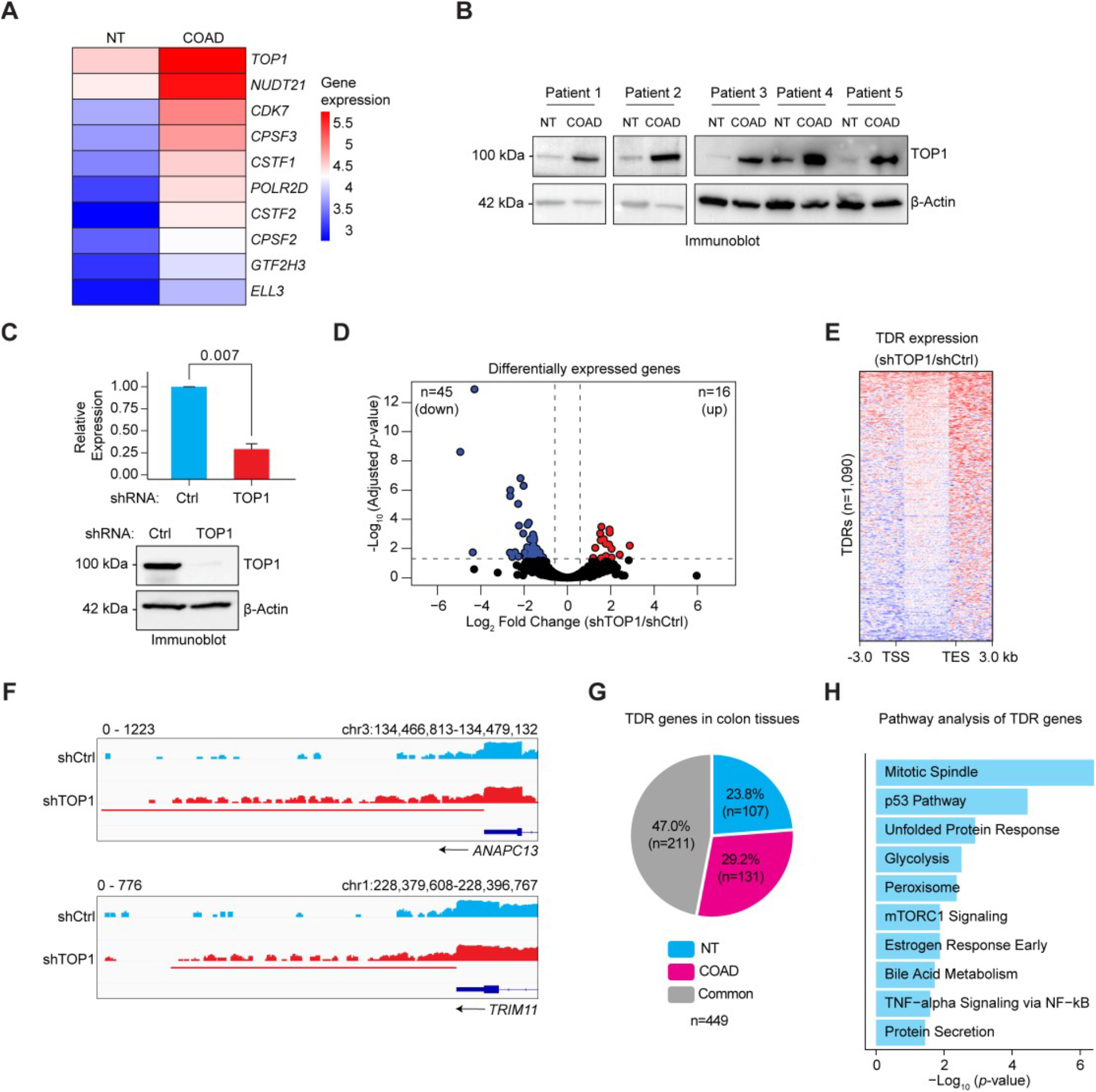
TOP1 regulates DoG RNA biogenesis in colorectal carcinoma. A, Heatmap showing the median expression of genes classified by GEPIA (*62*) as associating with RNAPII transcription and rank ordered by log2 fold change in expression as top ten most highly expressed genes in RNA-seq data from 275 CRCs relative to 349 NTs from TCGA and GTEx (*27, 34*). **B,** Immunoblot analysis of TOP1 and *β*-Actin from paired NT and COAD tissues of five individual patients. **C,** (top) qRT-PCR and (bottom) immunoblot analysis of SW480 cells stably expressing control shRNA (Ctrl) or TOP1 shRNA. Data represents the mean and s.e.m of three independent experiments. Statistical significance determined by two-tailed Student’s *t* test. (*p*-value = 0.007). **D,** Volcano plot of the differentially expressed RefSeq genes (log2 FC>0.58 or log2 FC<- 0.58, *q*-value< 0.05) in SW480 cells stably expressing control shRNA (Ctrl) or TOP1 shRNA. **E,** Heatmap of RNA-seq distribution spanning 3 kb upstream of the TSS to 3 kb downstream of the TES of the TRT genes (n = 1,090). The log2 ratio of RNA-seq signal (RPKM) is represented in TOP1 knockdown versus shCtrl SW480 cells. **F,** IGV tracks of RNA-seq signal (RPKM) at the *TRIM11* and *ANAPC13* loci in SW480 cells expressing Ctrl and TOP1 shRNA. The horizontal bar defines the DoG region determined by DoGFinder (*28*). **G,** Pie chart depicting the number of TDR genes that are DoG-producing genes in NT, COAD, or both (common) in patient tissues described in Fig. 2A. **H,** Top ten (*p*-value < 0.05) MSigDB pathways for TDR genes (n = 1,090). See also Fig. S4 and Table S5 and S6.

To identify whether TOP1 regulates the colon cancer transcriptome, we performed RNA-seq in SW480 colon cancer cells expressing short hairpin RNAs (shRNAs) against TOP1. As shown in Fig. 4C, relative to a non-targeting shRNA against LacZ (Ctrl), TOP1 shRNA markedly reduced TOP1 mRNA (top) and protein (bottom) levels. By comparing the transcriptomes of control and TOP1-depleted cells, we found that TOP1 significantly alters (< or > 1.5-fold change, *q*-value < 0.05) the expression of few (n = 61) genes (Fig. 4D and Table S5). This moderate effect of TOP1 on gene expression is consistent with a previous report revealing a similar number of genes regulated by TOP1 in breast cancer cells (*35*). Notably, however, we identified 1,090 genes that specifically produce DoG RNAs following TOP1 depletion as evidenced by the significant increase in transcription signal past the annotated TESs of these protein coding genes (Fig. 4E, Fig. S4B, and Table S6). The identification of DoG RNAs following TOP1 depletion is evidenced past the TESs of the *TRIM11* and *ANAPC13* gene loci (Fig. 4F). For brevity purposes, we refer to these DoG RNAs, whose expression is suppressed by TOP1 as the TOP1 DoG RNAs (TDRs). Consistent with TOP1 primarily regulating DoG RNA versus mRNA expression is the minimal change in RNA-seq signal spanning from the transcriptional start site (TSS) to the TES (Fig. 4E) and that only one of the TOP1-regulated DoG-producing genes was differentially expressed following TOP1 depletion (Fig. S4C). Additional analysis of the TDRs, revealed that these DoG RNAs are defined by significantly longer extension strength as compared to DoG RNAs in shCtrl cells, averaging 24.0 kb versus 22.5 kb, respectively (Fig. S4D). TOP1-dependent inhibition of DoG RNA production was further revealed by qRT-PCR that showed an increase in DoG RNAs, do*DAPK3* (2-fold), do*BRF1* (3-fold), and do*MAPK9* (2.8-fold) following TOP1 depletion (Fig. S4E). Moreover, TOP1 regulation of DoG RNAs, do*DAPK3,* do*BRF1,* and do*MAPK9* was further confirmed following TOP1 depletion using siRNAs directed against different regions of TOP1 mRNA (Fig. S4F) and following TOP1 knockdown in a second colon cancer HCT116 cell line (Fig. S4G). Altogether, these results show a role of TOP1 in block the expression of TDRs in colon cancer cells.

To further investigate the significance of TOP1-dependent inhibition of DoG RNA production in colon cancer, we examined whether TOP1 dysregulation in colon cancer tumors is associated with altered DoG RNA production. Using RNA-seq data from TCGA we identified 50 COAD tumors with variable TOP1 expression levels, 25 COAD tumors with low versus 25 tumors with high TOP1 expression levels (Fig. S4H, left). Using these 50 COAD tumors for DoG RNA identification, we identified 33% less DoG RNAs in the TOP1 high versus TOP1 low COAD tumors. Specifically, we identified 3,013 versus 1,011 DoG RNAs in the TOP1 low versus high COAD tumors (Fig. S4H, right), which is consistent with our findings that TOP1 depletion leads to increased induction of DoG RNAs. Moreover, comparative analysis of the DoG-producing host genes that support TDR production following TOP1 depletion in colon cancer cells revealed partial overlap (449 of 1,090) with the DoG-producing genes in NTs (n = 107, 24%), COAD tumors (n = 131, 29%) and those that are shared among the NTs and COAD tumors (n = 211, 47%) (Fig. 4G). Consistent with TOP1-regulated DoG RNAs overlapping with NT and COAD tumors is the identification that the host genes supporting production of DoG RNAs following TOP1 knockdown are enriched for both tumor promoting pathways (mTORC1 signaling, unfolded protein response, glycolysis, and mitotic spindle) and tumor suppressor pathways (p53 pathway and TNF-a signaling via NF-KB) (Fig. 4H). Taken together, these findings underscore the importance of TOP1 as an essential regulator of DoG RNA production in colon cancer.

## Paused RNAPII and catalytically inactive TOP1 accumulate at TES proximal regions

To investigate the mechanisms underlying the direct functions of TOP1 in preventing DoG RNA expression, we monitored the relationship between global TOP1 catalytic activity and the binding profiles of TOP1 and RNAPII. Using SW480 colon cancer cells, we performed chromatin immunoprecipitation followed by deep sequencing (ChIP-seq) for RNAPII and TOP1, and TOP1 catalytic activity was measured by employing TOP1- sequencing (TOP1-seq, Fig. 5A), a method for identifying only catalytically engaged TOP1 (TOP1cc) (*21*). To examine the relationship between TOP1cc, TOP1, and RNAPII profiles and transcription, we next parsed the TDR host genes and non-DoG producing genes in SW480 cells by using the lower and upper quartiles into three subsets, low (quartile 1), medium (quartile 2), and high (quartile 3) transcription levels as measured by RNA-seq (Fig. 5B). The TDR host genes are transcribed at high levels that are comparable to the transcription levels of the highly transcribed non-DoG producing genes (Fig. 5B). At the TSSs and gene bodies of TDR and highly transcribed non-DoG-producing genes, significantly higher levels of TOP1, RNAPII, and TOP1cc accumulation were observed relative to the low and medium transcribed non-DoG producing genes (Fig. 5C). The increased enrichment of TOP1, RNAPII, and TOP1cc at the TSSs and gene bodies are consistent with increased supercoiling that coincides with the significantly higher transcription levels observed at TDR and highly transcribed non-DoG producing genes (Fig. 5B). At the TES proximal regions, a second peak of TOP1 and RNAPII binding was observed that, similar to the TSS peak was significantly higher at the TDR host genes and the highly transcribed versus the low and medium transcribed non-DoG producing genes (Fig. 5C). Comparative analysis of the TOP1 ChIP-seq and TOP1- seq revealed significant overlap of TOP1 binding and catalytic activity at the TSSs of TDRs and all subsets of the non-DoG producing genes, respectively (Fig. 5C). In comparison, TOP1 catalytic activity and TOP1 binding are discordant in gene bodies and in TES proximal regions as evidenced by the negligible levels of TOP1cc versus the significant enrichment of TOP1 binding in these genomic regions (Fig. 5C). This data suggests a role of TOP1 occupancy instead TOP1 catalytic activity at these sites, which could be associated with the recruitment of key factors associated with DoG transcription. The high density of RNAPII binding that overlaps with TOP1 binding at the TSS and TES of TDR host genes is consistent with RNAPII pausing patterns. To examine whether the TDR host genes are largely occupied by paused RNAPII, we used our PRO-seq data in SW480 cells to calculate the pausing index (PI) for all DoG producing genes versus two subsets of non-DoG producing genes, the TDR host genes (yellow, Fig. 5B) and the highly transcribed non-DoG producing genes (green, Fig. 5B). Importantly, PRO-seq enables detection of nascent transcripts at a single nucleotide resolution that are specifically produced by paused or elongating polymerases versus stalled or arrested polymerases. As shown in Fig. 5D, we calculated the PI at the TSS (PITSS) by measuring the ratio of RNAPII density at the TSS (L1, −50 to +300 bp) relative to the RNAPII density in the gene body (L2, +300 bp to the annotated end of the genes). In addition, we measured the PI associated with 3’ gene ends (PITES) by dividing the normalized RNAPII density at the TES proximal region (L3, +500 to +1500 bp) by the RNAPII density downstream of the TES (L4, +1,500 bp to +10,000 bp). At the TSSs, a PITSS greater than 2 was identified, which is consistent with a strong promoter bias of paused RNAPII at all three subsets of genes (Fig. 5E). Moreover, we found that the median PITSS was significantly higher (n = 1,368) for the DoG producing genes as compared to the non-DoG producing genes, which were found to have comparable median PITSS, TDR host genes (x = 1,361) and the highly transcribed non-DoG producing genes (x = 2,279), respectively (Fig. 5E). Examination of the PITES further revealed an enrichment of paused RNAPII at all three subsets of genes (Fig. 5E), which is consistent with all three subsets of genes consisting of paused RNAPII patterns at the TSS and TES. Notably, and consistent with DoG production, we found that the median PITES associated with DoG-producing genes is significantly lower (X = 11) than the median PITES for the TDR host genes (X = 15) and the highly transcribed non-DoG producing genes (X = 75), respectively. Notably, the median PITES for the TDR host genes was found to be significantly lower than that of the non-DoG producing genes (Fig. 5E). Consistent with DoG-producing genes having the lowest median PITES is the finding that the PRO-seq signal remains at a comparable level at the TES proximal region and beyond (∼10 kb downstream) (Fig. S5). Similarly, and consistent with the lack of DoG production from the TDR and highly transcribed non-DoG producing genes is the finding that the accumulation of paused RNAPII at the TES proximal region is consistent with the sharp decrease in PRO-seq signal in the region downstream of the TES (Fig. S5). Taken together, these data demonstrate that TDR genes exhibit high levels of concordance between the binding profiles of TOP1 and paused RNAPII at the TSS, gene bodies, and TES proximal regions. Moreover, differences in the localization of TOP1 binding versus TOP1cc throughout protein coding genes suggests that TOP1 is likely to exhibit other, non-catalytic roles within gene bodies and TES proximal regions.

**Fig. 5.**
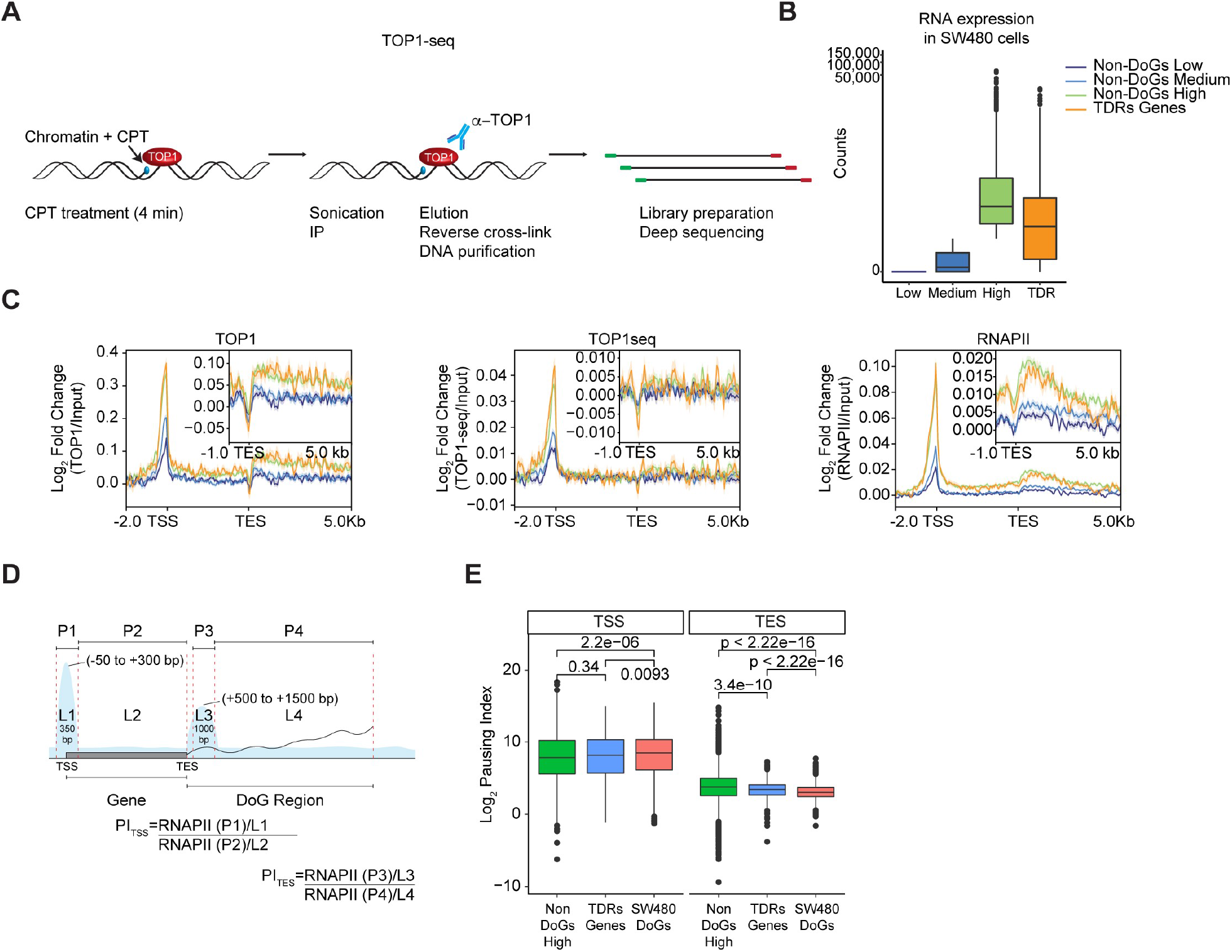
Paused RNAPII and catalytically inactive TOP1 accumulate at TES proximal regions. A, Schematic of TOP1-seq, a method to identify only catalytically engaged TOP1 in SW480 cells (*21*). **B,** Box plot showing the counts of TDRs and non-DoG-producing genes with low, medium, and high transcription in SW480 cells. The counts were grouped using the lower and upper quartiles into three groups, low (quartile 1), medium (quartile 2), and high (quartile 3) expression according to the Total RNA-seq reads. Boxplots enclose values between first and third quartiles, midlines show medians, and whiskers extend to data points within 1.5 the interquartile range from the box, outliers are shown. **C,** Metaplots of TOP1, TOP1cc, and RNAPII ChIP-seq signal at TDRs (n = 1,090) and non-DoG-producing genes with low, medium, and high transcription in SW480 cells. TOP-seq and ChIP-seq signal is represented as log2-transformed fold change of bins per million over Input and spans 2 kb upstream of the TSS to 5 kb downstream of the TES. A zoom-in spanning from −1 kb to +5 kb around the TES is shown. **D,** Illustration depicting the pause index determination for TDR genes. The TSS pause index is defined as the ratio of RNAPII reads at paused site 1 at the TSS (P1, spanning from −50 to +300 bp) over the RNAPII reads at gene body (P2, spanning from +300 to end of the gene). The TES pause index is defined as the ratio of RNAPII reads at paused site 2 on TES (P3, spanning from +500 to +1500 bp) over the RNAPII reads at DoG Region identified by DoGFinder (*28*) (P4, spanning from +1500 to end of the DoG defined by DoGFinder (*28*)). **E,** Box plot showing the TSS and TES pause index in highly expressed non-DoG genes (green), TDRs genes (blue), and SW480 DoG-producing genes (red) in SW480 cells. Statistical significance was determined by Wilcoxon rank-sum test. See also Fig. S5.

## TOP1 is essential for RNAPII termination at thousands of TDR host genes

Given the striking overlap of TOP1 and RNAPII binding profiles, we next wanted to examine whether TOP1-dependent regulation of DoG RNA expression is associated with defects in RNAPII behavior. First, we examined whether TOP1 and RNAPII form functional associations in SW480 cells. As shown in Fig. 6A, an antibody specifically recognizing TOP1 co-immunoprecipitated RNAPII from nuclear extracts prepared from SW480 cells. These results establish physiological associations between TOP1 and RNAPII that are consistent with the global overlap in their binding profiles (Fig. 5C).

**Fig. 6.**
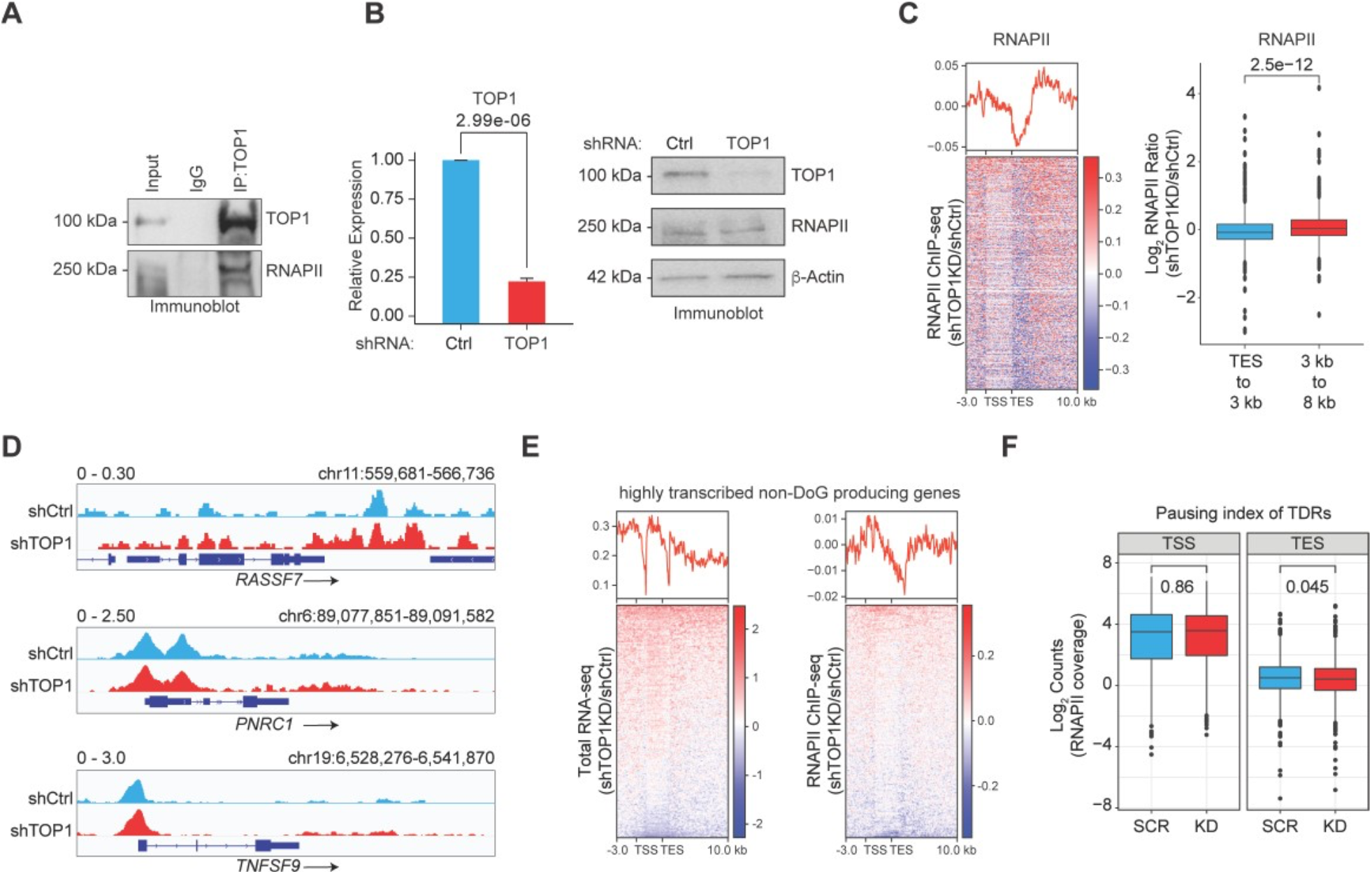
TOP1 interacts with RNAPII to regulate TDR genes. A, Co-immunoprecipitation with TOP1 and IgG (control) antibodies from SW480 nuclear extracts and immunoblot analysis of RNAPII and TOP1. A representative image is shown that is representative of three independent experiments. **B,** (left) qRT-PCR analysis of *TOP1* mRNA. Data represents the mean and s.e.m of three independent qRT-PCR experiments. Statistical significance was determined by two-tailed Student’s *t* test. *p*- value = 2.99e-06. (right) Immunoblot analysis of TOP1, RNAPII, and *β*-Actin in SW480 cells expressing Ctrl and TOP1 shRNA. A representative image is shown that is representative of three independent experiments. **C,** Heatmap of RNAPII ChIP-seq distribution spanning 3 kb upstream of the TSS to 10 kb downstream of the TES of the TDR genes (n = 1,090). The RNAPII ChIP-seq represent the log2 ratio of ChIP-seq signal in TOP1 knockdown over shCtrl SW480 cells. **D,** IGV tracks of RNAPII ChIP-seq signal at the *RASSF7, PNRC1* and *TNFSF9* loci in SW480 cells expressing Ctrl and TOP1 shRNA. **E,** Heatmaps of transcription and RNAPII ChIP-seq distribution spanning 3 kb upstream of the TSS to 10 kb downstream of the TES of the highly transcribed non-DoG genes in TOP1 KD (n = 2,325). The log2 ratio of RNA-seq signal (RPKM) is represented in TOP1 knockdown versus shCtrl SW480 cells. The RNAPII ChIP-seq represent the log2 ratio of ChIP-seq signal in TOP1 knockdown over shCtrl SW480 cells. **F,** Box plot showing the TSS and TES pause index on TDRs genes in SW480 cells expressing Ctrl and TOP1 shRNA. Statistical significance was determined by Wilcoxon rank-sum test.

We next examined whether TOP1 is directly regulating DoG RNA production by modulating RNAPII chromatin engagement. To examine this possibility, RNAPII ChIP-seq was performed following TOP1 shRNA-mediated knockdown, which significantly decreased *TOP1* mRNA and protein levels without affecting RNAPII protein levels (Fig. 6B). Inspection of the heatmap analyses (Fig. 6C, left) and individual (Fig. 6D) TDR host genes revealed decreased RNAPII binding at the TSS and TES proximal regions in TOP1 versus control knockdown conditions. The decrease in RNAPII levels at the TSS was not accompanied by a significant change in RNAPII binding within the coding regions of TDR genes, which is consistent with TOP1 depletion resulting in little to no change in the high expression levels of the TDR host genes (Fig. 4D). Notably, at the TES versus TSS, a greater decrease in RNAPII binding was observed, which was accompanied by a striking increase in RNAPII binding downstream of the TES proximal region (Fig. 6C, left). Quantitative analysis of RNAPII binding at the TES proximal region (TES to 3 kb downstream) and downstream of the TES proximal region (3 kb to 8 kb) revealed that RNAPII binding is significantly increased past the TES proximal regions (Fig. 6C, right). Moreover, and as expected, we did not observe a change in RNAPII binding in the DoG regions of the highly transcribed non-DoG producing genes following TOP1 depletion (Fig. 6E), which is consistent with these genes not producing DoG RNAs. As shown in Fig. 6C, the significant increase in RNAPII occupancy is consistent with the total RNA-seq (Fig. 4E) and PRO-seq (Fig. S3B) results that confirmed the presence of transcriptionally active RNAPII mapping to the regions past the TESs of TDR host genes following TOP1 depletion.

To determine whether increased RNAPII accumulation and induction of DoG RNA production following TOP1 depletion is due to the release of paused RNAPII into productive elongation, we calculated the PITES and PITSS using our RNAPII ChIP-seq data in control versus TOP1 knockdown conditions (Fig. 6F). Consistent with our PI calculations using PRO-seq data in SW480 cells, which revealed 1,044 genes with paused RNAPII (Fig. 5E, the RNAPII ChIP-seq data in shCtl cells revealed a comparable number of 865 genes with paused RNAPII at the TSS and 324 genes with paused RNAPII at the TES of TDR host genes (Fig. 6F). Analysis of the PITSS in control and TOP1 knockdown conditions revealed negligible changes in the PITSS following TOP1 depletion, which suggests that TOP1 does not regulate the release of paused RNAPII into the TDR gene bodies (Fig. 6F). Notably, however, the PITES at TDR genes was found to be significantly lower in TOP1 knockdown versus control cells (Fig. 6F), which is consistent with the observed increase in RNAPII levels that are detected downstream of the TES proximal region of the TDR genes following TOP1 knockdown (Fig. 6F). Collectively, these data reveal an essential and direct function of TOP1 in promoting termination at TDR host genes (Fig. 7).

**Fig. 7.**
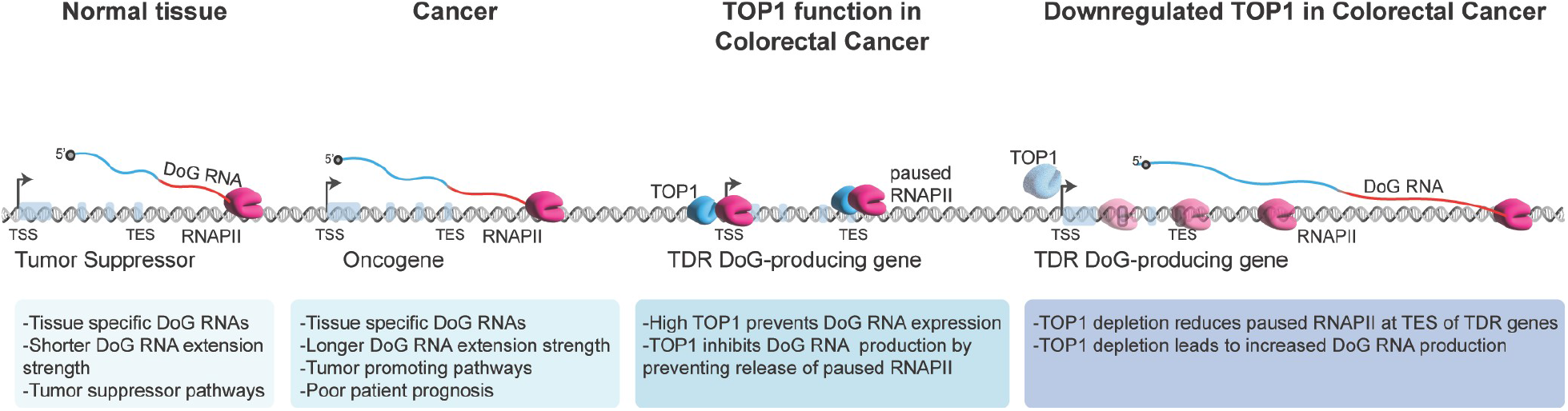
Model. Proposed model for how DoG RNAs re-shape normal and cancer transcriptomes. Comparative analyses of DoG RNA signatures in normal versus cancer tissues provides new insights into differentially expressed DoG RNAs in four major cancer types and revealed that DoG RNAs are largely tissue-specific. Dysregulated expression of DoG RNAs in breast and colon tumors are significantly correlated with poor patient survival. Consistent, DoG-producing host genes associated with the tumor-specific DoG RNAs encode for RNAs with known tumor-promoting functions. Mechanistically, high TOP1 expression in colorectal cancer prevents DoG RNA expression by preventing the release of paused RNAPII binding specifically at transcription end sites (TESs). Finally, TOP1 depletion promotes DoG RNA expression by lowering the levels of RNAPII pausing and promoting the release of RNAPII well beyond the ends of TDR host genes.

## Discussion

While noncoding RNAs (ncRNAs) make-up the vast majority of our transcriptome, our understanding of the noncoding genome remains limited. Among the emerging classes of ncRNAs are DoG RNAs that are induced in response to stress stimuli including osmotic stress (*3, 8*), viral infection (*5, 6*), and heat shock (*5–7, 36–38*). Several unsolved questions regarding DoG RNAs exist. For example, are DoG RNAs produced in normal cellular contexts and in human disease states including cancer? Can differential expression analyses of DoG RNAs in normal versus cancer tissues provide new glimpses into the biological differences between a healthy versus cancer cell? What makes certain protein coding genes in the genome prone to DoG RNA production, and related, are there specific transcriptional regulators that control DoG RNA expression? In our study, we provide new insights to address these questions by revealing the identification and classification of DoG RNAs and a new mechanism underlying their regulation and function in colon cancer. Collectively, our results presented here will foster future investigations of these ncRNAs in both normal cellular processes and human cancers.

To reach a deeper understanding of alterations in the cancer cell transcriptome, we sought to demonstrate the existence and cancer-specificity of DoG-producing host genes and their DoG RNAs. By employing an approach using TCGA RNA-sequencing data sets from matched tumors and NTs and established NGS pipelines (*28*), we reveal a comprehensive catalog of DoG RNA signatures that were previously unrecognized and importantly re-shape both normal and cancer transcriptomes. Comparative analysis of the DoG RNA signatures in NTs versus tumors that are of breast, colon, liver, and prostate origin revealed differentially expressed DoG RNAs in all four cancer types and that DoG RNAs are largely tissue-specific. Through deep transcriptomic profiling, we elucidated these DoG RNA signatures as potential biomarkers that inform cancer prognosis. Reasurringly, our approach independently recovered reproducible DoG RNA signatures in COAD tumors from TCGA and independent molecular profiling of a subset of paired NTs and COAD tumors from human patients and several colon cancer and normal colon epithelial cell lines. Regarding the functions of the elucidated DoG RNA signatures is our observation that the overlapping DoG RNA-producing genes in colon cancer encode for transcripts involved in tumor promoting functional pathways including MYC targets, mTORC1 signaling, G2-M checkpoint, and E2F targets. Comparatively, the functional annotations observed for DoG-producing genes specific to NTs revealed an enrichment of normal cellular and developmental processes. Remarkably, and consistent with DoG RNAs overlapping with biological hallmarks of tumorigenesis, we also found that dysregulated DoG RNAs in both breast and colon cancer are significantly correlated with poor patient survival. These findings are consistent with the growing noncoding genome and clinical significance of identifying and classifying new classes of ncRNAs for elucidating their potential relevance as biomarkers and therapeutic targets.

In addition to our observation of differential and tissue-specific expression of DoG RNAs in tumors versus normal tissues, our findings also reveal varying molecular features associated with the cancer-specific DoG RNAs. Specifically, we found that increased DoG number and extension strength define DoG RNAs that are specific to tumors versus NTs. Since DoG RNAs are continuous noncoding RNA extensions of their upstream protein coding genes, our study suggests the significance of increased DoG number and length in tumorigenesis. Consistent, our analysis revealed a clear pattern of increased DoG extension strength across all tumor samples from the various tissues that we surveyed. Remarkably, our findings also revealed that increased DoG number and extension strength are significantly associated with poor patient survival in both breast and colon cancer patients. Recent studies examining shifts in gene length support a paradigm for age-associated imbalances in gene-length (*39*). In addition to aging, length-associated transcriptomic imbalances have also been linked to Alzheimer’s disease (*40*). While varying length transcripts are enriched under specific biological contexts, future work is needed to determine causality of the varying length-associated transcripts including DoG RNAs in both normal development and disease states.

Our study reveals a previously unrecognized mechanism that involves a pivotal role for the essential enzyme TOP1 in preventing global DoG RNA production. This is evidenced by our identification of a thousand genes that “escape transcriptional repression” and support high levels of DoG production following TOP1 depletion. The power of TOP1 as a potent inhibitor of DoG production is consistent with the high prevalence of TOP1 dysregulation in cancer (*41*). Consistent, we identifed that TOP1 knockdown in colon cancer cell lines recapitulates our findings across multiple tumors in which low versus high TOP1 levels are associated with high versus low DoG RNA production. Also, consistent is the finding that TDR genes partially overlap with gene loci that specifically produce DoG RNAs in NTs and are largely associated with tumor suppressor pathways and that partially overlap with DoG-producting genes in COAD tumors that are known to promote tumorigenesis. Future investigations will be required to address whether TOP1 prevents DoG production to support tumorigenesis by inhibiting tumor suppressor pathways and/or by promoting pathways that enhancer cancer cell growth.

Importantly, TOP1 has long been viewed as a key determinant of gene control that is governed by its regulation of the preinitiation complex (PIC) (*14–20*) and RNAPII elongation (*21*). TOP1 regulates transcription initiation by enhancing the formation of an active TFIID-TFIIA complex (*15*) and by directly stimulating nucleosome disassembly and gene expression (*17*). TOP1 has also been shown to regulate DNA topology during elongation through reinforced TOP1 activity mediated by RNAPII (*21*). Importantly, our studies unveil that TOP1 is also a key regulator of gene control at the level of termination. Strikingly, the identification of increased DoG RNA production together with negligible changes in the levels of the DoG-producing genes upon TOP1 depletion is consistent with TOP1’s primary role in regulating termination. While RNAPII pausing has been shown to be a mechanism for regulating transcriptional activation, the extent to which it is involved in the control of gene expression and termination remains poorly understood.

Here we demonstrate a new role for TOP1 in directly influencing RNAPII pausing at the ends of thousands of genes. When this regulatory control is overcome by TOP1 depletion, RNAPII is released into productive elongation that, in turn gives rise to aberrant transcriptional activation downstream of thousands of protein coding genes. While promoter-proximal pausing of RNAPII is increasingly recognized as an important step in regulating gene expression, for at least a subset of genes (*42*), our studies represent a paradigm shift in our understanding of pausing defects in noncoding transcriptional regulation. Just as specific genes are defined by the accumulation of promoter proximal pausing including rapid response genes, we suggest that stably paused RNAPII shapes the regulation of an emerging class of DoG RNAs in colon cancer. Additional profiling analysis of histone modifications and transcriptional regulatory factors will continue to refine the roles for TOP1 and RNAPII pausing in regulating DoG RNAs.

We have uncovered a biological basis for DoG RNA signatures in human cancers through their annotation and characterization in reference to paired normal tissues for several major cancers that include colon, breast, liver, and prostate. Importantly, our study also unveils that these molecular vulnerabilities are under the control of a regulatory switch governed by an essential regulator of gene control, TOP1. By defining the previously unrecognized role of TOP1 in regulating termination, we now demonstrate the significance of TOP1 in regulating all stages of the RNAPII transcription cycle and further support the paradigm that DoG RNAs are increasingly identified as hallmarks of termination defects (*3, 5, 6, 8, 38, 43*).

## Materials and Methods

### Cell culture and treatments

Human colorectal adenocarcinoma DLD-1, SW480, HCT116 cells, normal colon HCoEpiC and FHC cells, and human embryonic kidney 293T (HEK293T) cells were purchased from the American Type Culture Collection (ATCC). DLD-1, SW480, HCT116, and HEK293T cell lines were grown in Dulbecco’s modified Eagle medium (DMEM, Gibco) supplemented with 10% fetal bovine serum (FBS, Gibco). The HCoEpiC cells were grown on poly-l-lysine coated plates with Colonic Epithelial Cell Medium (ScienCell) supplemented with 5 ml of colonic epithelial cell growth supplement (ScienCell) and 5 ml of penicillin/streptomycin solution (ScienCell). The FHC cells were grown in DMEM:F12 Medium (ATCC) supplemented with 25 mM HEPES, 10 ng/ml cholera toxin, 0.005 mg/ml insulin, 0.005 mg/ml transferrin, 100 ng/ml hydrocortisone, 20 ng/mL human recombinant EGF (Thermo Fisher PHG0311), and 10% fetal bovine serum. All cell lines were grown in a 37 °C incubator supplied with 5% CO2. All cell lines tested negative for mycoplasma contamination by PCR.

### RNA interference experiments using siRNA

SW480 cells were transfected with 100 nM non-targeting siRNA control (Ctrl) or TOP1 siRNA duplexes listed in Table S7 (Dharmacon) using Lipofectamine 3000 (Invitrogen) according to the manufacturer’s directions (Life Technologies). Cells were harvested 48 h post-transfection for immunoblot or RNA expression analyses.

### Lentivirus production and transduction

pLKO.1 TRC control and target shRNA plasmids were generated with annealed primers to knockdown TOP1. shRNA primers used in this study are listed in Table S7. For lentivirus production and transduction, 50– 60% confluent HEK293T cells were transfected with TRC control, target shRNAs, and packaging plasmids psPAX2 and pMD2.G using Lipofectamine 3000 (Invitrogen). Virus-containing medium was collected 48 and 72 h post transfection, filtered with a 0.45-μm pore size filter, and used for viral infection. SW480 and HCT116 cells were transduced with viral supernatants containing 8 μg ml^-1^ polybrene (Sigma-Aldrich). After 8h infection, virus-containing medium was removed and replaced with fresh medium. After 48 h, the cells were selected using puromycin at a final concentration of 1.5 μg ml^-1^ before harvesting the cells for qRT-PCR and immunoblot to confirm successful knockdown efficiency.

### Antibodies

Antibodies used for ChIP assays were obtained commercially as follows: anti-Topoisomerase I (abcam, ab219735, 2μg), anti-RNAPII (N-20) (Santa Cruz, Sc-899, 2μg), anti-Rpb1 NTD (D8L4Y) (Cell Signaling, #14958, 5 ul), anti-Spike-in antibody (Active Motif, 61686, 2μg), and anti-IgG (Cell Signaling, 2729S, 2μg). Antibodies used for immunoblotting are as follows: anti-*β*-Actin (Santa Cruz, sc47778, 1:1000 dilution), anti-Topoisomerase I (abcam, ab219735, 1:1000 dilution), anti-Rpb1 NTD (D8L4Y) (Cell Signaling, #14958, 1:1000 dilution).

### Colorectal carcinoma tumor analysis

Nine colorectal carcinoma tumors and their corresponding paired normal colon tissue were obtained from the Department of Pathology of Northwestern University, Feinberg School of Medicine following informed consent from patients. Samples were dissected by pathologists and frozen in liquid nitrogen for molecular analysis. Briefly, for RNA extraction, 1 mL of TRIzol Reagent (Invitrogen) was added per 50 mg of tissue into a homogenizing tube (Precellys). For protein extraction, 1 mL of RIPA lysis buffer (10mM Tris-HCl pH 8.0, 1mM EDTA, 0.5mM EGTA, 1% Triton X-100, 0.1% NaDoc, 0.1% SDS, 140mM NaCl) was added per 50 mg of tissue into a homogenizing tube (Precellys). Tissues were homogenized with the Bertin Technologies power homogenizer for 10 cycles, 20 seconds per cycle at speed setting #2. Following homogenization, samples were centrifuged at 12,000 x g for 10 min. at 4°C, and the cleared lysate was used for RNA purification or immunoblotting as described in detail below.

### Western blotting

Protein samples were denatured at 95°C for 5 min, separated by SDS-PAGE, and transferred to PVDF membranes using the iBlot2 gel transfer device (Invitrogen). The membranes were blocked in 3% milk and probed with the indicated antibodies. Reactive bands were detected by Pierce™ ECL Plus (Thermo Scientific Pierce) or SuperSignal™ West Femto (Thermo Scientific Pierce) and visualized using the Odyssey Fc Imaging System (LI-COR Biosciences) or the ChemiDoc Imaging system (Bio-Rad Laboratories).

### RNA Purification and qRT-PCR

Total RNA was extracted with TRIzol reagent (Invitrogen) from 9 paired patient tissues (CRCs and NTs), SW480, HCT116, DLD-1, HCoEpiC, and FHC cells, SW480 and HCT116 cells stably expressing Ctrl or TOP1 shRNAs or SW480 cells transiently expressing control, TOP1 siRNAs following manufacturer’s instructions. RNA (100 ng) was used for cDNA synthesis using the ProtoScript II First Strand cDNA Synthesis Kit (NEB) with random hexamers. PCR reactions were performed using SYBR Green PCR Master Mix (Applied Biosystems) on an Applied Biosystems QuantStudio3 real-time PCR system in triplicate using samples from at least three independent cell harvests. The specificity of amplification was confirmed by melting curve analysis. The relative levels of DoG and mRNA expression were calculated using the ΔΔCt method normalized to *β*-Actin (tumor samples) or GAPDH (cell lines). The expression levels in TOP1 knockdown are relative to the Ctrl knockdown. Primers for qRT-PCR are listed in Table S7.

### RNA-sequencing

Total RNA was extracted from 5 paired patient tissues (COAD and NTs), HCT116, SW480, FHC cells, and from SW480 cells stably expressing Ctrl or TOP1 shRNAs using TRIzol reagent (Invitrogen) according to the manufacturer’s instructions. Genomic DNA was removed by Turbo DNase treatment (Invitrogen). Total RNA-seq (ribo-depleted) libraries for all samples were generated and sequenced by Admera Health. Briefly, isolated RNA sample quality was assessed by High Sensitivity RNA Tapestation (Agilent Technologies Inc., California, USA) and quantified by Qubit 2.0 RNA HS assay (ThermoFisher, Massachusetts, USA). Libraries were constructed with KAPA™ RNA HyperPrep with RiboErase (Roche, Indiana, USA) and performed based on manufacturer’s recommendations. Final library quantity was measured by KAPA SYBR® FAST qPCR and library quality evaluated by TapeStation D1000 ScreenTape (Agilent Technologies, CA, USA). The final library size was about 430 bp with an insert size of about 200bp. Illumina® 8-nt dual-indices were used. Equimolar pooling of libraries was performed based on QC values and sequenced on an Illumina® NovaSeq S4 (Illumina, California, USA) with a read length configuration of 150 PE for 50 M PE reads per sample (25M in each direction).

### TOP1-sequencing

The TOP1-seq methodology was adapted from a previously established protocol (*21*). Briefly, 20-24×10^6^ SW480 cells were treated with 10 µM CPT for 4 min. Cells were immediately washed with ice-cold 1X PBS, scraped, resuspended in lysis buffer (20 mM Tris-HCl pH 7.5, 300 mM NaCl, 2 mM EDTA, 0.5% NP40, 1% Triton X-100, 1 mM PMSF, PICs), and incubated on ice for 15 min. Cell resuspensions were dounced ten times using an ice-cold glass homogenizer. Nuclei were collected by centrifugation and resuspended in shearing buffer (0.1% SDS, 0.5% N-lauroylsarcosine, 1% Triton X-100, 10 mM Tris-HCl pH 8.1, 100 mM NaCl, 1 mM EDTA, 1 mM PMSF, PICs). Chromatin was fragmented using a bioruptor Pico sonicator (Diagenode) to an average size of 200-600 bp and cleared chromatin was used for immunoprecipitation overnight at 4°C with protein A Dynabeads that were pre-coupled in 0.5% BSA with IgG or TOP1 antibodies. The immunocomplex-bound beads were washed eight times in wash buffer (50 mM HEPES-KOH pH 7.6, 500 mM LiCl, 1 mM EDTA, 1% NP40, 0.7% sodium deoxycholate, 1 mM PMSF, PICs), twice in 1X TE, and eluted in elution buffer (50 mM Tris-HCl pH 8.0, 10 mM EDTA, 1% SDS). Immunoprecipitated DNA samples were purified using UPrep® Spin Columns (Genesee Scientific) and quantified with a Qubit 4.0 fluorometer (Invitrogen). We used 10 ng of DNA from two biological replicates to generate sequencing libraries using KAPA Hyper Prep Kit (KAPA Biosystems) according to the manufacturer’s instructions. Briefly, TOP1-seq were subjected to end-repair, adaptor ligation followed by indexed PCR. Libraries were size-selected for average size of 300 bp. Sequencing was performed on an Illumina® NovaSeq 6000 (Illumina, California, USA) with a read length configuration of 75 SE per sample.

### Precision nuclear run-on sequencing

PRO-seq was performed following established protocol (*44*). Nuclei from SW480 cells that were isolated and snap-frozen in storage buffer (10 mM Tris-HCl pH 8.0, 25% glycerol, 5 mM MgCl2, 0.1 mM EDTA and 5 mM DTT). Next, 10×10^6^ nuclei 100 µL storage buffer were run on by the addition of 100 µL pre-warmed 2X-reaction buffer [10 mM Tris-HCl pH 8.0, 300 mM KCl, 5 mM MgCl2, 1% Sarkosyl, 1 mM DTT, 0.8 U μL^-1^ RNaseOUT (ThermoFisher Scientific), 250 µM ATP, 250 µM GTP, 50 µM biotin-11-CTP (PerkinElmer), and 50 µM biotin-11-UTP (PerkinElmer)] for 3 min at 37°C. Reactions were stopped by the addition of 500 µL TRIzol LS (Invitrogen) and purified following the manufacturer’s instructions. RNAs were heat-denatured at 65°C for 40 sec, placed on ice, and fragmented with 0.2 N NaOH on ice for 10 min. After pH adjustment with 1 M Tris-HCl pH 6.8, buffer exchange using P-30 columns (Bio-Rad Laboratories) was performed. Biotinylated RNAs were enriched with 30 µL of pre-washed Dynabeads MyOne Streptavidin C1 (Life technologies) in binding buffer (10 mM Tris-HCl pH 7.4, 300 mM NaCl and 0.1% Triton X-100, 0.8 U μL^-1^ RNaseOUT) and by gentle rotation at room temperature for 30 min. Beads were subsequently washed twice with ice-cold high-salt wash buffer (50 mM Tris-HCl pH 7.4, 2 M NaCl and 0.5% Triton X-100), twice with binding buffer, once with low-salt wash buffer (5 mM Tris-HCl pH 7.4 and 0.1% Triton X-100), and eluted with TRIzol LS (Invitrogen). The 3’ RNA adaptor (VRA3: 5’- GAUCGUCGGACUGUAGAACUCUGAAC-/inverted dT/-3’) was ligated to the Eluted RNAs and a second biotin RNA enrichment was performed as described above. Next, the 5’ end of RNAs were enzymatically modified with RNA 5’ Pyrophosphohydrolase (RppH; NEB) and hydroxyl repair was performed using T4 Polynucleotide Kinase (PNK; NEB). Purified RNAs were subsequently ligated with the 5’ RNA adaptor (RA5: 5’- CCUUGGCACCCGAGAAUUCCA-3’). A third biotin RNA enrichment was performed prior to reverse transcription of the eluted RNAs. Libraries from two biological replicates were amplified for 13 cycles, size selected for 140–350 bp, and single-end sequenced on Illumina HiSeq 4000 (75bp).

### Chromatin Immunoprecipitation

TOP1 and RNAPII ChIP assays were performed using 24-36×10^6^ SW480 cells and RNAPII ChIP assays were performed using the same number of SW480 cells stably expressing control or TOP1 shRNAs. The cells were cross- linked with 6 mM disuccinimidyl glutarate (DSG; ProteoChem) for 30 min followed by crosslinking with 1% formaldehyde for an additional 10 min at room temperature and quenched with 125 mM glycine (Fisher Scientific) for 5 min. Cell pellets were resuspended in lysis buffer (20 mM Tris-HCl pH 7.5, 300 mM NaCl, 2 mM EDTA, 0.5% NP40, 1% Triton X-100, 1 mM PMSF, PICs) and incubated on ice for 30 min. Cell resuspensions were next dounced ten times in a prechilled glass homogenizer. Nuclei were collected and resuspended in shearing buffer (0.1% SDS, 0.5% N-lauroylsarcosine, 1% Triton X-100, 10 mM Tris-HCl pH 8.1, 100 mM NaCl, 1 mM EDTA, 1 mM PMSF, PICs). Chromatin was fragmented using a bioruptor Pico (Diagenode) or an E220 Focused-Ultrasonicator (Covaris) to an average size of 200-600 bp and the cleared chromatin was used for immunoprecipitation overnight at 4°C with Protein A Dynabeads that were pre-coupled in 0.5% BSA with IgG, TOP1, RNAPII, and Spike-in antibodies. The immunocomplex-bound beads were washed eight times in wash buffer (50 mM HEPES-KOH pH 7.6, 500 mM LiCl, 1 mM EDTA, 1% NP40, 0.7% sodium deoxycholate, 1 mM PMSF, PICs), twice in 1X TE, and eluted in elution buffer (50 mM Tris-HCl pH 8.0, 10 mM EDTA, 1% SDS). Crosslinks were reversed at 65°C overnight and the DNA samples were purified using UPrep® Spin Columns (Genesee Scientific). ChIP-seq libraries for all samples were generated and sequenced by Admera Health. Briefly, immunoprecipitated DNA was quantified with Qubit 2.0 DNA HS Assay (ThermoFisher, Massachusetts, USA) and quality assessed by Tapestation genomic DNA Assay (Agilent Technologies, California, USA). Library preparation was performed using KAPA Hyper Prep (Roche, Basel, Switzerland) following the manufacturer’s recommendations. All samples were subjected to end-repair, adaptor ligation followed by indexed PCR using Illumina® 8-nt dual-indices. Library quality and quantity were assessed with Qubit 2.0 DNA HS Assay and Tapestation High Sensitivity D1000 Assay (Agilent Technologies, California, USA). Final libraries were quantified using QuantStudio ® 5 System (Applied Biosystems, California, USA) prior to equimolar pooling based on qPCR QC values. Sequencing was performed on an Illumina® NovaSeq (Illumina, California, USA) with a read length configuration of 150 PE for 40M PE reads 20M in each direction) per sample.

### Co-Immunoprecipitation

Co-Immunoprecipitation was performed following established protocol (*45*). Briefly, 12×10^6^ cells were lysed in lysis buffer (10 mM Tris-HCl pH 7.4, 10 mM KCl, 1.5 mM MgCl2, 12% Sucrose, 10% Glycerol, 0.2% Triton X-100, 0.5 mM DTT, 1 mM PMSF, PICs, phosphatase inhibitors), then fractionated in sucrose cushion (10 mM Tris-HCl pH 7.4, 10 mM KCl, 1.5 mM MgCl2, 30% Sucrose, 0.5 mM DTT). Nuclei were stored in freeze buffer (10 mM Tris-HCl pH 7.4, 10 mM KCl, 1.5 mM MgCl2, 40% Glycerol, 0.5 mM DTT) at –80 C. Chromatin DNA was digested in Chromatin digestion buffer (20 mM Tris-HCl pH 7.4, 150 mM NaCl, 1.5 mM MgCl2, 10% Glycerol, 0.05% NP40, 0.5 mM DTT, 1 mM PMSF, PICs, phosphatase inhibitors, 250 U/ml Benzonase). After centrifugation, the supernatant was collected as fraction 1, while proteins were further extracted from the pellet in Chromatin-2 buffer (20 mM Tris-HCl pH 7.4, 500 mM NaCl, 1.5 mM MgCl2, 10% Glycerol, 0.05% NP40, 0.5 mM DTT, 1 mM PMSF, PICs, phosphatase inhibitors, 3 mM EDTA) as fraction 2. NaCl concentration in fraction 2 was adjusted to 150 mM, then combined with fraction 1. TOP1 immunoprecipitation was performed using anti-Topoisomerase I (abcam, ab219735, 2μg), conjugated to Protein A Dynabeads for 24h. The immunocomplex-bound beads were washed four times in wash buffer (20 mM Tris-HCl pH 7.4, 225 mM NaCl, 1.5 mM MgCl2, 10% Glycerol, 1.5 mM EDTA, 0.05% NP40, 0.5 mM DTT), immunoprecipitated proteins were eluted in Laemmli SDS sample buffer before immunoblotting analysis for TOP1 and RNAPII.

### Total RNA-seq and PRO-seq analysis

The quality control of raw data was evaluated with FastQC and MultiQC (*46*). The reads were filtered for quality using Trim Galore. Paired-end raw reads were mapped to the hg38 human genome using HISAT2 v2.2.1 (*47*) with default parameters with specific strand information. The gene expression was determined with featureCounts using RefSeq gene annotation and the next parameters -t exon -g gene_id (*48*). The gene counts were used to determine the differential gene expression with DESeq2 v1.34.0 (*49*) and q-value<0.05 and a log2 fold change of 0.58. Volcano plots were generated using ggplot2. The gene set enrichment analysis was carried out using enrichr (*29*). Terms with a p- value < 0.05 were deemed statistically significant. The boxplot, barplot, dotplot, venn diagram, and statistical analysis were performed in the ggplot2 package of R (*50*).

### DoG RNA Identification

BAM files from our total RNA-seq analysis aligned to the hg38 human genome were sorted and indexed with SAMtools v1.6 (*51*). DoG RNAs were identified using DoGFinder (*28*) with the following parameters, Get_DoGs -minDoGLen 5000 -minDoGCov 0.6 -w 200 -mode F using the protein-coding genes RefSeq annotation. Downsampling of the following RNA-sequencing data sets (i) NT and COAD samples for each of the 5 individual patients, (ii) cell lines including the SW480, HCT116 and FHC, and (iii) the shTOP1 and shCtrl cell lines was performed to reduce the differences in sequencing depth by using the Pre_Process function in DoGFinder. The DoG coordinates provided by DoGFinder were used to identify the counts coverage using Deeptools (*52*) that was used to generate bigwig files for visualization. RPKM normalization method was used and the log2 ratio between samples was calculated (COAD samples over NT samples, SW480 and HCT116 over FHC cells, and shTOP1 over shCtrl cells). Box plots were generated using ggplot2 (*50*). The adjacent genes were identified by using the *closest* function available in bedtools v2.30.0 (*53*).

### TCGA data analysis

To identify the DoG signature in cancer we use RNA-seq data available in the TCGA database (*27, 34*). RNA-seq files of breast invasive carcinoma (BRCA), colon adenocarcinoma (COAD), liver hepatocellular carcinoma (LIHC), and prostate adenocarcinoma (PRAD) patients from TGCA were downloaded from the Genomic Data Commons with the dbGaP accession phs000178.v10.p8. For this study, only paired-end format files were used. The BAM files were converted to fastq using SAMtools (*51*) and aligned to the hg38 human genome using HISAT2 with default parameters. The BAM files were sorted and indexed with SAMtools (*51*). DoG RNAs were identified with DoGFinder (*28*) by using Get_DoGs -minDoGLen 5000 -minDoGCov 0.6 -w 200 -mode F using the RefSeq annotation. TOP1 levels in COAD patients were determined by using the UCSC Xena platform (*54*) and ranked by increasing values.

Kaplan–Meier curves were generated by using the overall survival time in days of patients from TCGA downloaded from UCSC Xena platform (*54*). The median of DoG number and extension strength for BRCA, COAD, LIHC, and PRAD was determined by using the best available cut-off value from KMplot. The significance was determined by using a Log-Rank test.

To identify the differentially expressed genes in COAD versus NTs that are associated with RNAPII-dependent transcriptional regulation, we used the RNA-seq data available for 275 COAD and 349 NTs from TCGA and GTEx databases (*27, 34*). We analyzed the gene expression of 161 annotated genes classified according to the GO term, transcription by RNA polymerase II (GO: 0006366) (*31–33*). Then, we sorted the genes by log2FC (COAD/NT) in gene expression and subsequently sorted the top 10 genes found to be highly expressed in the COAD versus NTs, which were used to generate the heatmap by utilizing the ggplot2 package of R (*50*).

### ChIP-seq and TOP1-seq Analysis

The quality control of raw data was evaluated with FastQC and MultiQC (*46*) on Galaxy platform (*55*). The low-quality reads were removed using Trim Galore. Sequencing reads were mapped to the hg38 human genome using Bowtie2 v2.4.2 (*56*) and default parameters. SAMtools v1.11 (*51*) was used for filtering the mapping quality and the duplicate reads. Only uniquely mapped reads were used for further analysis. Spike-in normalization was performed by mapping the reads to the Drosophila melanogaster genome (dm6) using Bowtie2 (*56*). Peak calling and broad TOP1cc regions were identified using MACS2 v2.1.1(*57*) and a cutoff p-value of 0.05 for TOP1cc peaks, p<0.005 for TOP1 peaks, and q<0.05 for RNAPII peaks. Deeptools v3.3.2 (*52*) was used to generate normalized bigwig files by combining reads from two replicas. The bins per million mapped normalized reads were used and the log2 ratio between the ChIP-seq signal over Input was calculated. The bigwig files were visualized in IGV (*58*). Peak annotation was carried out using ChIPseeker (*59*).

### TSS and TES Pausing Index

To calculate the TSS pausing index, we used the previously described method (*60*). Deeptools (*52*) was used to analyze the coverage of RNAPII counts (PRO-seq or RNAPII ChIP-seq) at promoters of TDRs Genes (−50 bp to +300 bp around TSS) and the gene body (+300 downstream to the annotated gene end). The pausing index was calculated by using the following formula (Figure 5D):

PITSS = Read Counts at TSS/Length1/Read Counts at Gene body/Length2 Where “Length1” is the length of the promoter region (350 bp) and “Length2” is the length of the gene body region (+300 downstream of the TSS to the annotated gene end). A pausing index of 2 was used to define the paused TDRs Genes.

To define the TES pausing index we used a similar method as was used for calculating the TSS pausing index. Deeptools was used to analyze the coverage of RNAPII counts (PRO-seq or RNAPII ChIP-seq) at +500 bp to +1,500 bp around TES and +10,000 downstream to the annotated gene end for non-DoG-producing genes. For TDRs and DoGs in SW480 cells, we used the region spanning from +1,500 downstream to the annotated gene end to the end of the DoG defined by DoGFinder. The region +500 bp to +1,500 bp around the TES was defined according to RNAPII enrichment at TES regions (Figure 5C). The Figure 5D shows an illustration of the TSS and TES pausing indices.

## Data availability

RNA-seq, PRO-seq, ChIP-seq, and TOP1-seq, data have been deposited at GEO and are publicly available. Accession number: GSE202408. This paper analyzes existing, publicly available TCGA data. Accession number: phs000178.v10.p8.

## Acknowledgments

We thank the Lauberth lab members, Ali Shilatifard, and Marc Morgan for helpful discussions, and Marta Iwanaszko for advice on NGS data analysis. This research was supported in part through the computational resources and staff contributions provided by the Genomics Compute Cluster, which is jointly supported by the Feinberg School of Medicine, the Center for Genetic Medicine, Feinberg’s Department of Biochemistry and Molecular Genetics, the Office of the Provost, the Office for Research, and Northwestern Information Technology. The Genomics Compute Cluster is part of Quest, Northwestern University’s high-performance computing facility, with the purpose to advance research in genomics.

## Author information

Conceptualization, S.M.L, J.X., and A.L.P.A; methodology, most of the experiments were performed by J.X., A.L.P.A., and N.M.; computational analysis, J.X., A.L.P.A; software, A.L.P.A and J.X., investigation, J.X., A.L.P.A., N.M, resources, N.M., A.L.P.A., J.X., G.Y., S.D., R.S.; writing-review and editing, S.M.L, J.X., A.L.P.A., G.Y; funding acquisition, S.M.L. All authors critically reviewed the manuscript and approved the final version.

## Competing Interests

The authors declare no competing interests.

## Notes

### Competing Interest Statement

The authors have declared no competing interest.

