## Supplemental Figures for "Downstream-of-gene (DoG) transcripts contribute to an imbalance in the cancer cell transcriptome"

This file includes Supplementary figures 1-5

**Supplementary Figures**

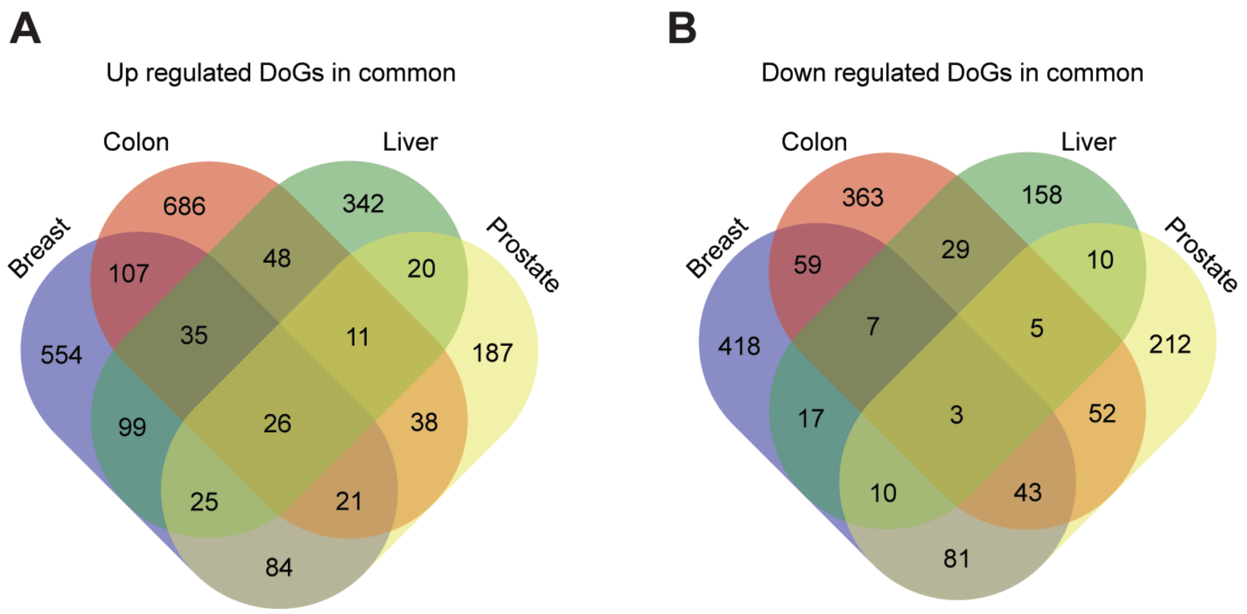


**Fig. S1. Common and specific DoG RNAs in breast, colon, liver, and prostate cancer.**

**A,** Venn diagram showing the overlap of up-regulated DoG RNAs in breast, colon, liver, and prostate tumors. **B,** Venn diagram showing the overlap of down-regulated DoG RNAs in breast, colon, liver, and prostate tumors.

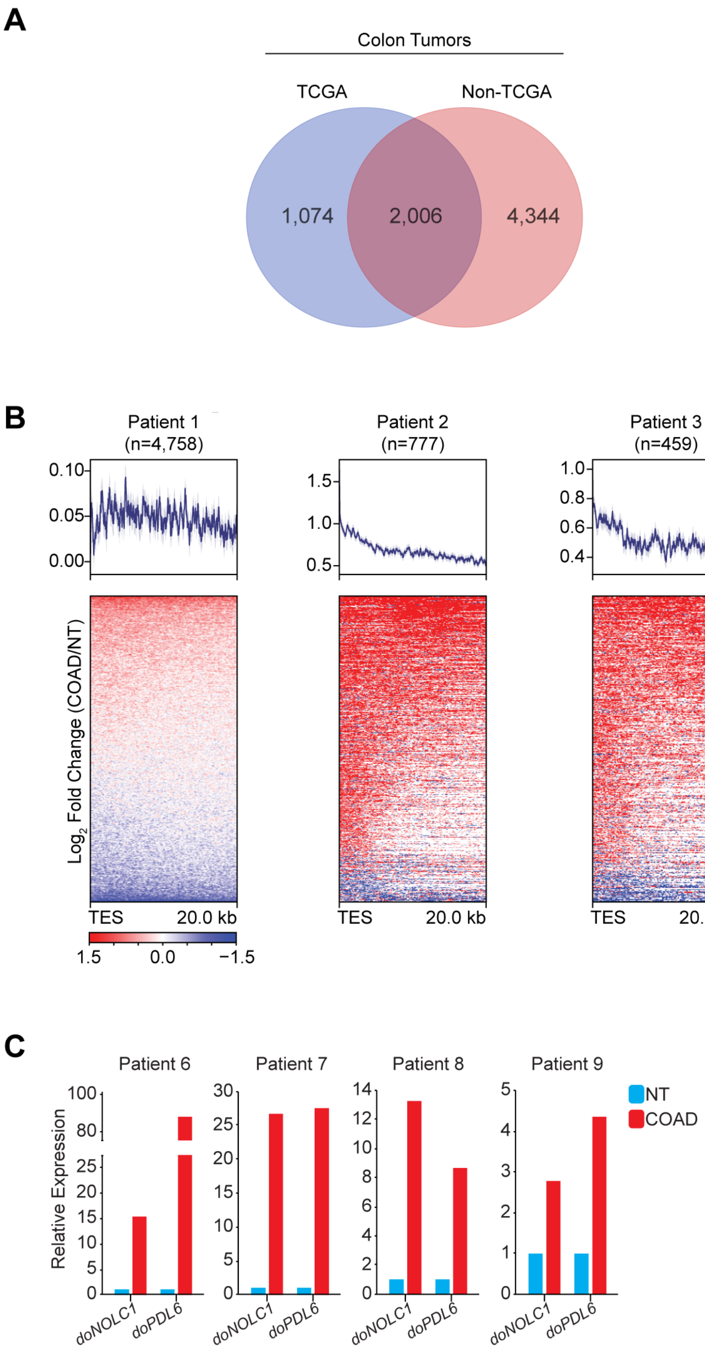


**Fig. S2. DoG RNA production is prevalent in patients with colorectal carcinoma.**

**A,** Venn diagram showing the overlap of DoG RNAs in the TCGA versus non-TCGA COADs (left) and TCGA versus non-TCGA normal colon tissues from this study (right). **B,** Heatmaps of the $log_2$-transformed fold change in RNA-seq signal (COAD/normal tissue, five patients) in Reads Per Kilobase Million (RPKM) spanning from the TES to

20.0 kb downstream of all annotated genes. **C,** qRT-PCR analysis of do*NOLC1* and do*PDL6* DoG expression in paired normal and tumor tissues from four additional colorectal cancer patients.

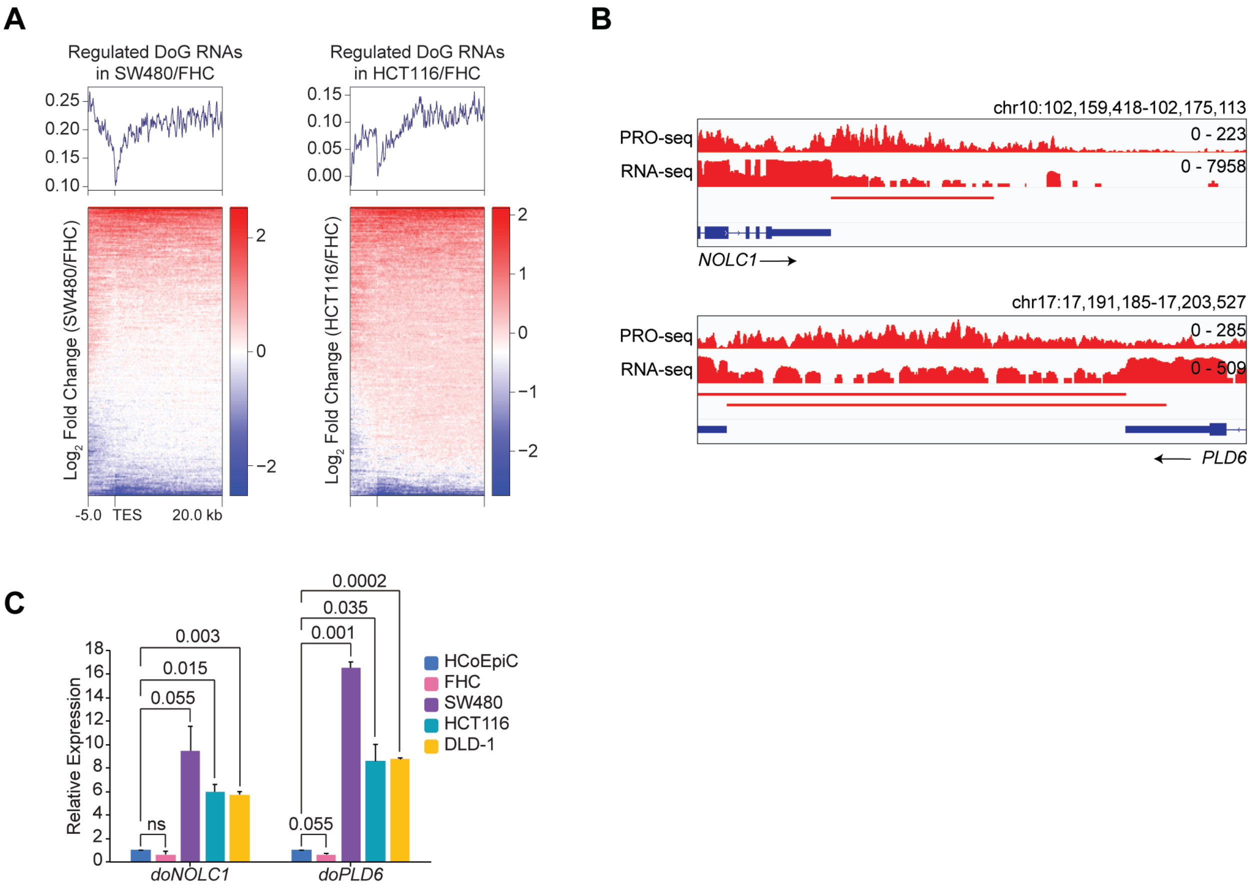


**Fig. S3. DoG RNAs in colorectal carcinoma cell lines.**

**A,** Heatmaps of the $log_2$-transformed fold change in RNA-seq signal (SW480/FHC and HCT116/FHC) in Reads Per Kilobase Million (RPKM) spanning from the TES to 20.0 kb downstream of all annotated genes. **B,** IGV tracks of PRO-seq and RNA-seq signal (RPKM) in SW480 cells of *NOLC1* and *PDL6* loci spanning the TES to the predicted end of the DoG as defined by DoGFinder(*28*). The horizontal bar defines the DoG regions identified by DoGFinder (*28*). **C,** qRT-PCR analysis of *doNOLC1* and *doPLD6* DoG RNA expression in HCoEpiC, FHC, SW480, HCT116, and DLD-1 cells. Data represents the mean and s.e.m. of two experiments that are representative of three independent experiments. Statistical significance was determined by two-tailed Student's *t* test. Significant *p*-values are shown, and ns (not significant).

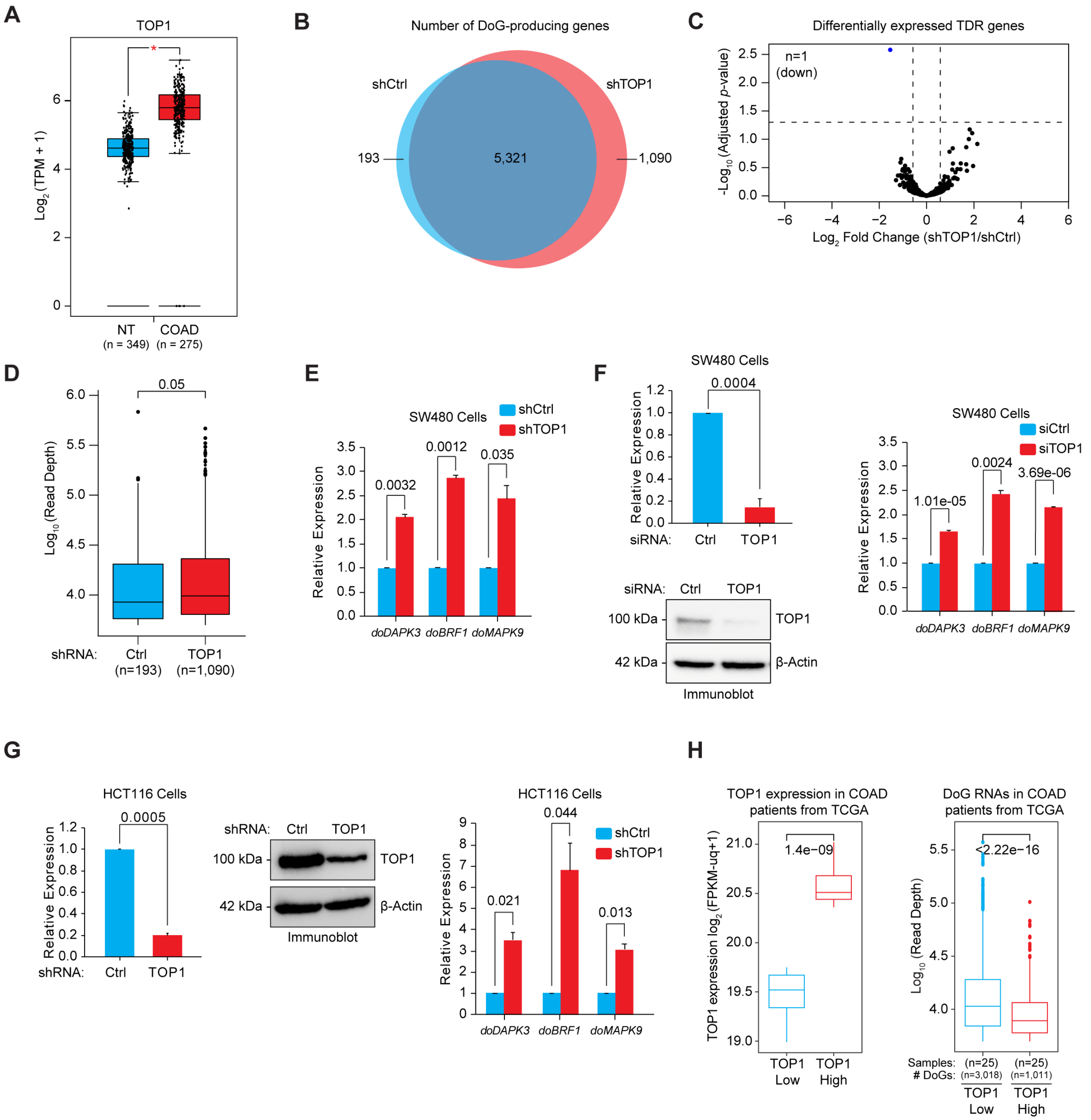


**Fig. S4. Depletion of TOP1 leads to potent DoG RNA production.**

**A,** Box with jitter plot for TOP1 RNA-seq levels in normal and colorectal adenocarcinoma tissues determined with GTEx and TCGA data from GEPIA server (*62*). Statistical significance was determined by one-way ANOVA test (**p*-value < 0.05). **B,** Venn diagram showing the overlap of DoG-producing genes in SW480 cells expressing Ctrl and TOP1 shRNA. **C,** Volcano plot showing differentially expressed TDR genes ($\log_2$ FC > 0.58, $\log_2$ FC < -0.58, *q*-value < 0.05) in SW480 cells expressing Ctrl and TOP1 shRNA. **D,** Extension strength of DoG RNAs identified by DoGFinder (*28*) in SW480 cells expressing Ctrl or TOP1 shRNA. The extension strength is shown in $\log_{10}$ scale. Boxplots enclose values between first and third quartiles, midlines show medians, and whiskers extend to data points within 1.5 the interquartile range from the box, outliers are shown. Statistical significance was determined by Wilcoxon rank-sum test. *p*-value=0.05. **E,** qRT-PCR analysis of the DoG RNA expression for *doDAPK3, doBRF1, and doMAPK9* in SW480 cells expressing Ctrl and TOP1 shRNA. Expression levels are relative to Ctrl shRNA. Data represents the mean and s.e.m. of two that are representative of three independent replicates. Statistical significance was determined by two-tailed Student's *t* test. *p*-values for *doDAPK3*=0.0032, *doBRF1*=0.0012, and *doMAPK9* =0.035. **F,** (left) qRT-PCR of TOP1 mRNA and TOP1 immunoblot analysis of SW480 cells transfected with Ctrl and TOP1 siRNA (n=3). β-Actin was used as loading control. Statistical significance was determined by two-tailed Student's *t* test. *p*-value=0.0004. (right) qRT-PCR analysis of the DoG RNAs for *doDAPK3, doBRF1,* and *doMAPK9* in SW480 cells transfected with Ctrl and TOP1 siRNA. Expression levels are relative to Ctrl siRNA. Data represents the mean and s.e.m. of two that are representative of three independent replicates. Statistical significance was determined by two-tailed Student's *t* test. *p*-values for *doDAPK3*=1.01e-05, *doBRF1*=0.0024, and *doMAPK9*=3.69e-06. **G,** (left) qRT-PCR and immunoblot (center) analysis of TOP1 in HCT116 cells transfected with Ctrl and TOP1 siRNA (n=3). β-Actin was used as loading control. Statistical significance was determined by two-tailed Student's *t* test. *p*-value=0.0005. (right) qRT-PCR analysis of the DoG RNAs for *doDAPK3*, *doBRF1*, and *doMAPK9* in HCT116 cells transfected with Ctrl and TOP1 siRNA. Expression levels are relative to Ctrl siRNA. Data represents the mean and s.e.m. of two that are representative of three independent replicates. Statistical significance was determined by two-tailed Student's *t* test. *p*-values for *doDAPK3*= 0.021, *doBRF1*=0.044, and *doMAPK9*=0.013. **H,** (left) COAD patients with low and high expression of TOP1. The boxplot shows TOP1 expression levels in the low and high subsets that were determined by parsing 25 patients with the lowest and 25 patients with the highest TOP1 levels from RNA-seq data of 100 TCGA CRC patients. Boxplots enclose values between first and third quartiles, midlines show medians, and whiskers extend to data points within 1.5 the interquartile range from the box, outliers are shown. Statistical significance was determined by Wilcoxon rank-sum test. (right) Number and extension strength of DoG RNAs identified in COAD patients with low and high expression of TOP1 from TCGA. Statistical significance was determined by Wilcoxon rank-sum test.

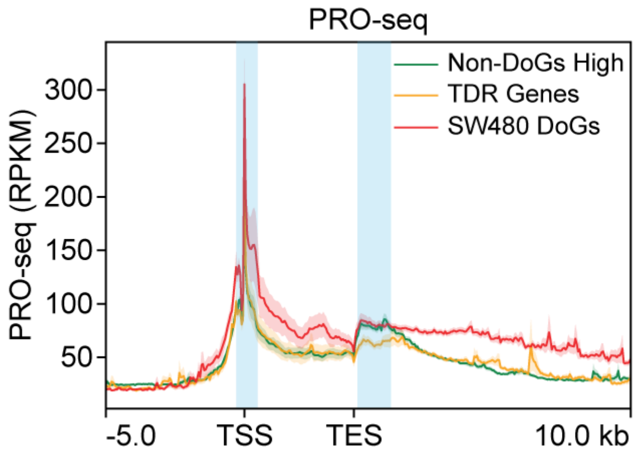


**Fig. S5. Accumulation of paused RNAPII at the TES proximal region in DoG-producing genes in SW480 cells.**

Metaplot of PRO-seq signal at highly expressed non-DoG genes (green), TDRs genes (yellow), and SW480 DoG-producing genes (red) in SW480 cells. PRO-seq distribution (RPKM) spanning 5 kb upstream of the TSS to 10 kb downstream of the TES of the genes is represented.
